# Context-dependent 3D genome regulation by cohesin and related factors

**DOI:** 10.1101/2022.05.24.493188

**Authors:** Ryuichiro Nakato, Toyonori Sakata, Jiankang Wang, Luis Augusto Eijy Nagai, Gina Miku Oba, Masashige Bando, Katsuhiko Shirahige

**Affiliations:** Laboratory of Computational Genomics, Institute for Quantitative Biosciences, University of Tokyo, 1-1-1 Yayoi, Bunkyo-Ku, Tokyo 113-0032, Japan; Laboratory of Genome Structure and Function, Institute for Quantitative Biosciences, University of Tokyo, 1-1-1 Yayoi, Bunkyo-Ku, Tokyo 113-0032, Japan; Karolinska Institutet, Department of Biosciences and Nutrition, Biomedicum, Quarter A6, 171 77, Stockholm, Sweden

**Keywords:** Computational genomics, 3D genome, transcriptome, epigenome, cohesin, CTCF, NIPBL

## Abstract

Cohesin plays vital roles in chromatin folding and gene expression regulation, cooperating with such factors as cohesin loaders, unloaders, acetyltransferase, and the insulation factor CTCF. Although various models of regulation have been proposed (e.g., loop extrusion), how cohesin and related factors collectively or individually regulate the hierarchical chromatin structure and gene expression remains unclear. In this study, we have depleted cohesin and related factors and then conducted a comprehensive evaluation of the resulting 3D genome, transcriptome and epigenome data. We observed substantial variation in depletion effects among factors at topologically associating domain (TAD) boundaries and on interTAD interactions, which were partly related to epigenomic status. Gene expression changes were highly correlated with direct cohesin binding and gain of TAD boundaries than with the loss of boundaries. Our results suggested that cohesin positively regulates gene expression, whereas other mechanisms (e.g., cohesin turnover and acetylation) add to the diversity of this pattern of dysregulation. Moreover, cohesin was broadly enriched in active compartment A, but not in compartment B, which were retained even after CTCF depletion. Our rich dataset and the subsequent data-driven analysis support the context-specific regulation of chromatin folding by cohesin and related factors.

## INTRODUCTION

The cohesin complex is crucial for gene transcription and chromatin folding in mammalian cells (Merkenschlager and Nora 2016; van Ruiten and Rowland 2018). Cohesin colocalizes with the CCCTC-binding factor CTCF to function as an insulator (Wendt et al. 2008), whereas a small proportion of cohesin binds the genome independently of CTCF, regulating gene expression with tissue-specific transcription factors (Schmidt et al. 2010; Faure et al. 2012). At least a subset of CTCF-independent cohesin mediates chromatin interactions between enhancer and promoter sites of active genes with mediator complexes (Kagey et al. 2010). Cohesin also participates in transcription elongation machinery that interacts with RNA polymerase II (Pol2) (Izumi et al. 2015). Mutations in the cohesin loader NIPBL (∼60%) and in cohesin subunits (∼10%) have been found in the human developmental disorder Cornelia de Lange syndrome (CdLS) (Kline et al. 2018).

Recent studies using whole-genome chromatin-conformation capture (Hi-C) uncovered a hierarchical three-dimensional (3D) genome structure regulated by cohesin and its related factors. Chromosomes are spatially segregated into active “compartment A” and inactive “compartment B” (Lieberman-Aiden et al. 2009). At a finer scale, chromosomes are folded into topologically associating domains (TADs), the boundaries of which are strongly enriched for cohesin and CTCF (Dixon et al. 2012). TADs can be nested, and interactions between TADs (interTAD interactions) are more rare than those within TADs (intraTAD interactions) (Bonev and Cavalli 2016). The depletion of cohesin or NIPBL causes a dramatic loss of TADs and chromatin loops (Rao et al. 2017; Schwarzer et al. 2017), whereas CTCF depletion affects TAD boundaries and loops more locally (Nora et al. 2017). These observations can be explained by the “loop extrusion” model, in which cohesin extrudes chromatin until it encounters CTCF, resulting in the formation of TADs (Fudenberg et al. 2016; Davidson et al. 2019; Kim et al. 2019). This model can also explain depletion effects of cohesin unloading factors (WAPL, PDS5A and PDS5B), which prevent the release of cohesin from DNA and cause loop extension, resulting in the appearance of longer loops than usual (Haarhuis et al. 2017; Wutz et al. 2017). Importantly, such extended loops are rare but also occur in wild-type cells (Allahyar et al. 2018), suggesting that CTCF boundaries are not absolute and the dynamics of TADs/loop formation may depend on the amount of cohesin on chromatin, which is balanced by continuous loading and unloading (turnover).

Despite these extensive efforts, the detailed mechanism of the hierarchical chromosome organization and the functional relationships involved in regulating transcription are still unclear (Sikorska and Sexton 2020). Extensive loss of TADs/loops and loop extension have a limited impact on gene expression and does not cause the spread of histone modifications (Haarhuis et al. 2017; Nora et al. 2017; Rao et al. 2017; Schwarzer et al. 2017; Ghavi-Helm et al. 2019). A dCas9-mediated insertion of boundary sequence was insufficient for creating TAD boundaries *de novo* (Bonev et al. 2017). Moreover, cohesin and CTCF also localize within TADs without forming boundaries. These results suggest a more complicated set of rules for chromatin structure formation and gene expression regulation by cohesin and related factors than the current models. Although each cohesin-related factor has been studied using different cell lines, a study to explore how cohesin and its related factors collectively or individually regulate chromatin folding, gene expression and the epigenome is needed.

Here we conducted a large-scale *in situ* Hi-C analysis after depletion of a variety of cohesin-related factors, with multiple replicates (31 samples, 14 billion paired-end reads in total). Combined with transcriptome and epigenome marks data, we comprehensively evaluated the similarities and differences in the resulting effects after depletion of individual factors. The resulting extensive dataset and subsequent analysis provide new insights into the context-specific roles of cohesin-related factors on gene expression and chromatin folding.

## RESULTS

### Datasets

Here we used human retinal pigment epithelial (RPE) cells to avoid the effect of aneuploidy or other genomic rearrangements (Figure 1A). We depleted cohesin (Rad21), cohesin loaders (NIPBL and Mau2), cohesin unloaders (WAPL, PDS5A and PDS5B), boundary element (CTCF) and cohesin acetyltransferase (ESCO1) and also carried out two sets of co-depletions (Rad21 and NIPBL, PDS5A and B). We confirmed that the depletion efficiencies were sufficient for all samples (Figure 1A) and that the majority of asynchronous cells were in G1 phase (Figure S1A). We used a 72-h treatment with short interfering RNA (siRNA) for most samples, but we also explored the effect of different treatment times (24, 48 and 120 h, Figure 1B). We also generated a sample that had been treated with the BET bromodomain inhibitor JQ1 because the bromodomain protein BRD4 is reported to interact with NIPBL and to be mutated in CdLS (Olley et al. 2018). Using these samples, we prepared *in situ* Hi-C, RNA-seq and spike-in ChIP-seq data (Tables S1–3). In the spike-in ChIP-seq, we observed that 60–80% of the peaks in control cells were lost after siRNA (Figure S1B).

**Figure 1.**
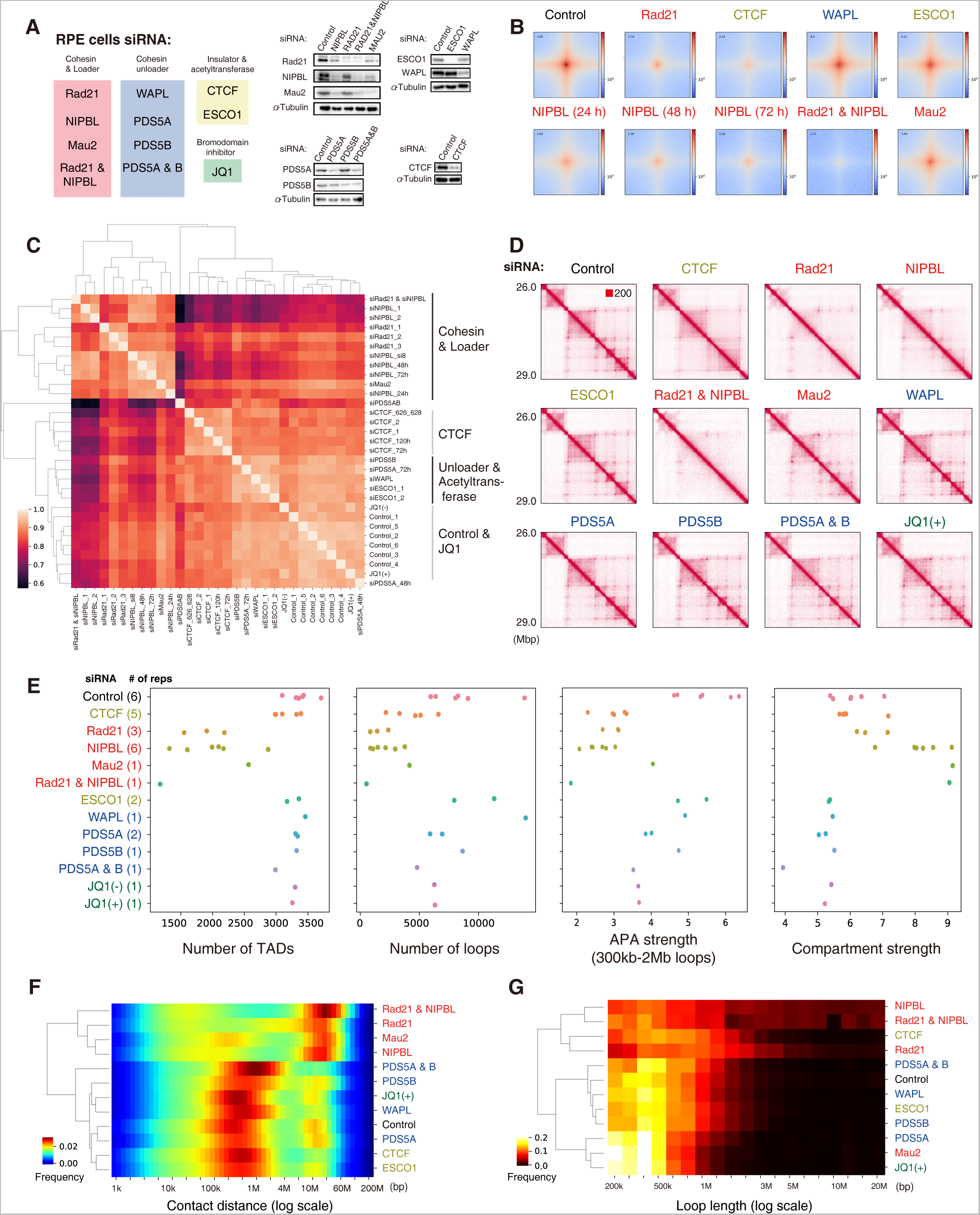
Comparative Hi-C analysis to explore the variation of depletion effects. **(A)** Summary of protein targets (left) and immunoblots after 72 h of siRNA (right). **(B)** Aggregate peak analysis (APA) of loops. **(C)** Correlation heatmap among samples based on stratum-adjusted correlation coefficients. **(D)** Normalized Hi-C matrix of a representative chromosomal region (chromosome 21, 26.0–29.0 Mb). **(E)** The number of TADs, loops, APA strength (observed/expected ratio of the center) and compartment strength (from saddle plots, see also Figure S2B). The dots indicate the Hi-C samples (including different treatment times). **(F)** Relative contact probability for the likelihood of a contact at increasing length scales (averaged by replicates). **(G)** Loop length distribution.

We evaluate the overall similarity among our Hi-C samples and found that the depletion effects can be categorized into four groups that correspond to the siRNA target (Figure 1C): cohesin and loaders, CTCF, cohesin unloaders and acetyltransferase, and control and JQ1. The exception was PDS5A and B co-depletion, which showed less correlation with NIPBL depletion (siNIPBL). Cohesin unloader and acetyltransferase depletion showed a milder effect on chromosome structure as compared with the cohesin loading and localization at CTCF sites. Having confirmed the sufficient similarity among replicates, we merged all replicates into a single deep Hi-C dataset for control, siRad21, siNIPBL (except for the 24-h treatment), siCTCF and siESCO1, resulting in at most 3 billion reads, for further analysis of these depletions.

### Comparative Hi-C analysis reveals diverse depletion effects on chromatin folding

We first analyzed Hi-C data and compared the depletion effects on TADs and loops. We observed a dramatic loss of TADs and loops after siRad21 and siNIPBL (Figures 1D and 1E), consistent with the previous studies (Rao et al. 2017; Schwarzer et al. 2017). Co-depletion of Rad21 and NIPBL showed a more severe effect. Mau2 depletion showed a milder effect than siNIPBL, possibly because some amount of cohesin can be loaded without Mau2 (Haarhuis et al. 2017). Although CTCF depletion strongly affected loops, it had a limited effect on TAD numbers and intraTAD interactions as compared with cohesin depletion (Figures 1D and 1E), indicating the function of CTCF as a boundary element (Nora et al. 2017; Hansen 2020). Most loops in the control samples anchored convergent CTCF motif sites as reported (Vietri Rudan et al. 2015), which was slightly violated after siWAPL, siPDS5AB and siCTCF (Figure S1C).

Compartmentalization can be uncoupled from TAD formation, which is strengthened by the depletion of cohesin and loaders (Haarhuis et al. 2017; Rao et al. 2017; Schwarzer et al. 2017) but not of CTCF (Nora et al. 2017). We observed a similar tendency in our data, as indicated by the “plaid pattern” (Figure S2A) and quantitative compartment strength estimated by a saddle plot (Figures 1E and S2B). siMau2 showed stronger compartmentalization than siRad21, in contrast to its milder effect on TADs and loops, suggesting the importance of cohesin loaders for compartmentalization. This tendency was also indicated by the relative contact frequency of mapped reads (Figure 1F). The depletion of cohesin and loaders diminished interactions at a length consistent with TADs (∼1 Mb), whereas the long-range interactions corresponding to the compartment (∼10 Mb) drastically increased. In contrast, depletion of CTCF, WAPL, PDS5B, PDS5AB or ESCO1 decreased long-range interactions, suggesting weakened compartmentalization. The depletion of PDS5A alone did not show a clear tendency. Lastly, we did not observe prominent compartment switching among any of the samples (Figure S2C).

We next explored the loop length distribution that showed a distinct tendency from the relative contact probability (Figure 1G). After siRad21, most short loops were depleted, and the distribution peaked at a longer length (∼10 Mbp) than did the control (∼500 kbp). siCTCF showed a similar but less drastic effect. After siNIPBL and after NIPBL and Rad21 co-depletion, a dramatic loss of short loops was observed, whereas a small number of long-range interactions appeared (> 5 Mbp, possibly due to the cohesin-independent long-range loops (Rao et al. 2017)). In contrast, siMau2 resulted in highly depleted long loops (∼1 Mbp), and the distribution then peaked at a shorter length than the control (∼400 kbp), which was similar to the effect of siPDS5A and JQ1. Based on the loop extrusion model, it was likely that shorter loops were retained under the mild loss of cohesin after siMau2. After depletion of PDS5B, PDS5A and B or WAPL, the peak distribution increased slightly relative to control samples (∼500 kbp), consistent with their function as cohesin unloaders. siESCO1 also caused the appearance of longer loops, similar to the effect of siPDS5B.

Additionally, we investigated the allele-specific depletion effect on chromosome X. Whereas the active chromosome X (Xa) forms the typical chromosome structure, inactive chromosome X (Xi) is partitioned into two megadomains, the boundary between which was affected by depletion of cohesin (Wang et al. 2018; Kriz et al. 2021). Although our data did not show an explicit disruption of the megadomain boundary in Xi, possibly due to incomplete siRNA depletion, we did observe a difference in depletion effects between Xi and Xa (Figure S3). Xi showed a “coarser” plaid pattern than Xa, which was strengthened by siRad21 and siNIPBL. siCTCF showed an asymmetric tendency of interaction frequency between the megadomain boundary and the two megadomains (black arrows), whereas there was no similarly clear chromosome-wide pattern in Xa. In addition, the interaction within the smaller megadomain was less affected in all samples. In summary, our Hi-C analysis showed consistent tendencies with previous studies, confirming its reliability, and provided multiple new findings of diverse depletion effects on chromatin folding.

### Gene expression changes were correlated with direct cohesin binding

Next, we explored the depletion effect on gene expression. We detected 2,000–7,000 differentially expressed genes (DEGs) for each sample (false discovery rate [FDR] < 0.01; Figures 2A and S4A). We selected the top-ranked 1,000 DEGs from all samples and merged them into a single DEG list (4,240 genes in total). Pairwise comparisons showed extensive overlap of DEGs between cohesin and loaders and between individual unloaders (Figure 2A). Interestingly, siNIPBL was more similar to siPDS5B than siRad21 and siMau2, suggesting DEGs from dysregulation of cohesin turnover. siCTCF and JQ1 showed less correlation with the others, suggesting their distinct roles for gene expression regulation.

**Figure 2.**
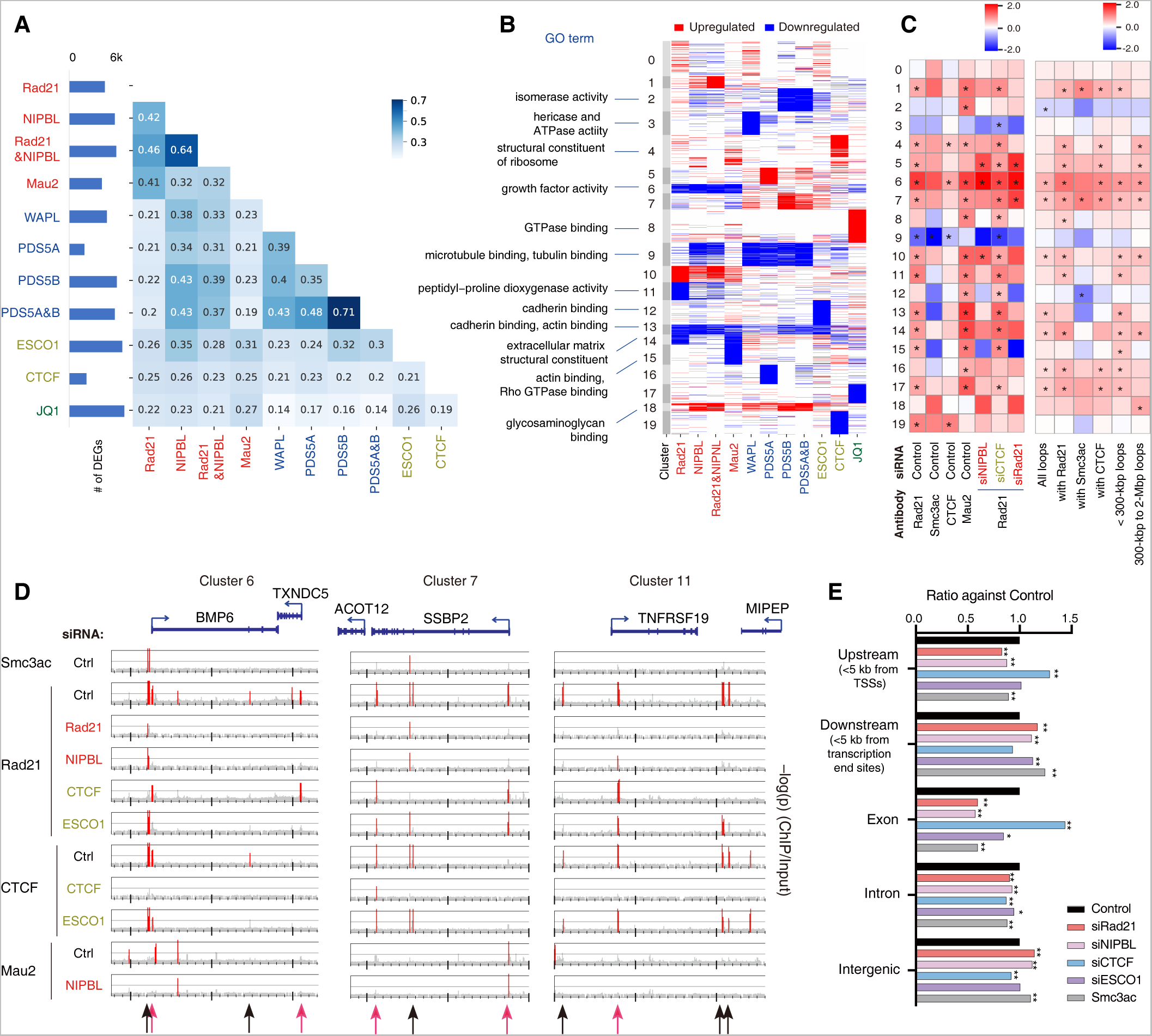
Comparative analysis of siRNA effects on the transcriptome and epigenome. **(A)** Left: Number of DEGs for each siRNA treatment (FDR < 0.01). Right: Correlation heatmap based on the Simpson index (showing overlap among top-ranked 1,000 DEGs) for all sample pairs. **(B)** Clustering of all top-ranked DEGs (rows) and samples (columns) based on the overlap of up- and downregulated genes. The significant GO terms are also shown (left). See Figure S5 for the full list of GO terms. **(C)** Log-scale relative enrichment of ChIP-seq peaks (left) and Hi-C loops (right) at the TSSs of DEGs in the clusters corresponding to **(B)** against all DEGs. *p < 0.01, the permutation test (n = 1,000). **(D)** ChIP-seq distribution (–log_10_(p), 100-bp bin) among top-ranked DEGs (*BMP6, SSBP2* and *TNFRSF19*). The significantly enriched regions (p < 10^−4^) are highlighted in red. Red and black arrows (bottom) indicate CTCF-independent and -dependent Rad21 peaks, respectively. **(E)** The relative enrichment of Rad21 peaks after depletions and Smc3ac around genes, as compared with Rad21 (control). *: p < 0.001; **: p < 0.0001, Fisher exact test against control.

To identify the pattern of expression dysregulation, we applied k-means clustering (k = 20) based on the overlap of up- and downregulated genes among siRNA treatments (Figure 2B and Table S4). For example, clusters 6 and 10 represent down- and upregulated genes after cohesin and loader depletion, respectively. Gene ontology (GO) analysis suggested that cluster 6 was mainly enriched in “growth factor activity,” consistent with slower growth under such depletions (Waizenegger et al. 2000). Clusters 9 and 18 contained down- and upregulated genes after NIPBL and unloader depletions, respectively. These DEGs were not observed after siRad21 and therefore would be correlated with cohesin turnover. Their GO terms were correlated with fundamental functions related to the cytoskeleton and extracellular matrix. These diverse expression patterns suggested multiple roles for cohesin-related factors in gene expression regulation.

We next examined the enrichment of ChIP-seq peaks and Hi-C loops at transcription start sites (TSSs) of the DEGs (Figure 2C). Most clusters were enriched for Rad21 and Mau2 peaks, suggesting that expression dysregulation of these clusters was caused by loss of Rad21 and Mau2 binding to each gene, rather than by region-wide effects caused by TAD disruption. We also found that the siRad21 DEGs were less likely to be located around disrupted TADs compared with non-differential boundaries (Figure S4B), suggesting little correlation between TAD disruption and gene expression dysregulation after siRad21. The exception was downregulated genes after siWAPL (cluster 3 and 9), which were independent of cohesin binding, implying the indirect or unrelated regulation relative to cohesin. In addition, loops mediated by acetylated cohesin (Smc3ac) were enriched in upregulated DEGs associated with siNIPBL and siPDS5B (clusters 1, 7, 18), whereas clusters not enriched for Smc3ac (12, 13, 15) were downregulated. Considering that acetylated cohesin sites are more stable and thus were more persistent even under siNIPBL and siRad21 as compared with non-acetylated sites (Figure 2C), this result suggests the necessity of cohesin at TSSs for gene expression.

Figure 2D shows the ChIP-seq distribution around several top-ranked DEG loci, each of which have cohesin peaks around their TSSs. Remarkably, Rad21 peaks around TSSs were lost after siNIPBL, whereas they remained after siCTCF (red arrows), suggesting that cohesin binds at TSSs in a more CTCF-independent manner. In contrast, Rad21 peaks in other regions were lost after siCTCF (black arrows). We confirmed that this tendency was genome-wide (Figure 2E). siNIPBL significantly depleted cohesin peaks at upstream and exon regions, whereas siCTCF affected intron and intergenic regions. In summary, our result suggested that cohesin positively regulates gene expression via direct binding at TSSs, whereas other mechanisms (e.g., turnover and acetylation) add to the diversity of this pattern of dysregulation.

### Quantitative classification of insulation levels reveals diversity among boundaries

To further study the depletion effects on chromatin folding, we calculated a multi-scale insulation score (Crane et al. 2015) (Figures 3A and 3B). We found various patterns of insulation perturbation at TAD boundaries: (i) boundaries weakened by siRad21 and siNIPBL but not by siCTCF (cohesin-dependent), (ii) boundaries strengthened by siNIPBL and siRad21 (cohesin-separated), (iii) boundaries depleted by siRad21, siNIPBL and siCTCF (all-dependent) and (iv) boundaries that were barely affected by any siRNA (robust). To quantify the observed patterns across the genome, we classified all boundaries into six types based on insulation score (Figure 3C and Table S5). In this classification, over half of the boundaries were annotated “robust,” indicating their stability in the presence of reduced amounts of a targeted protein. Most of the rest were classified as weakened boundaries after siRNA. Depletion of unloader proteins did not show an explicit perturbation, which is consistent with their minimal influence on the number of TADs (Figure 1E). Regarding the comparison with compartments, CTCF-dependent and cohesin-separated boundaries occurred more frequently between compartments A and B, whereas cohesin-dependent ones occurred less frequently (Figure 3D). This result suggests that cohesin has a role in connecting neighboring TADs (Schwarzer et al. 2017), especially those from compartments A and B, whereas CTCF is involved in partitions within compartment A and, to a lesser extent, compartment B.

**Figure 3.**
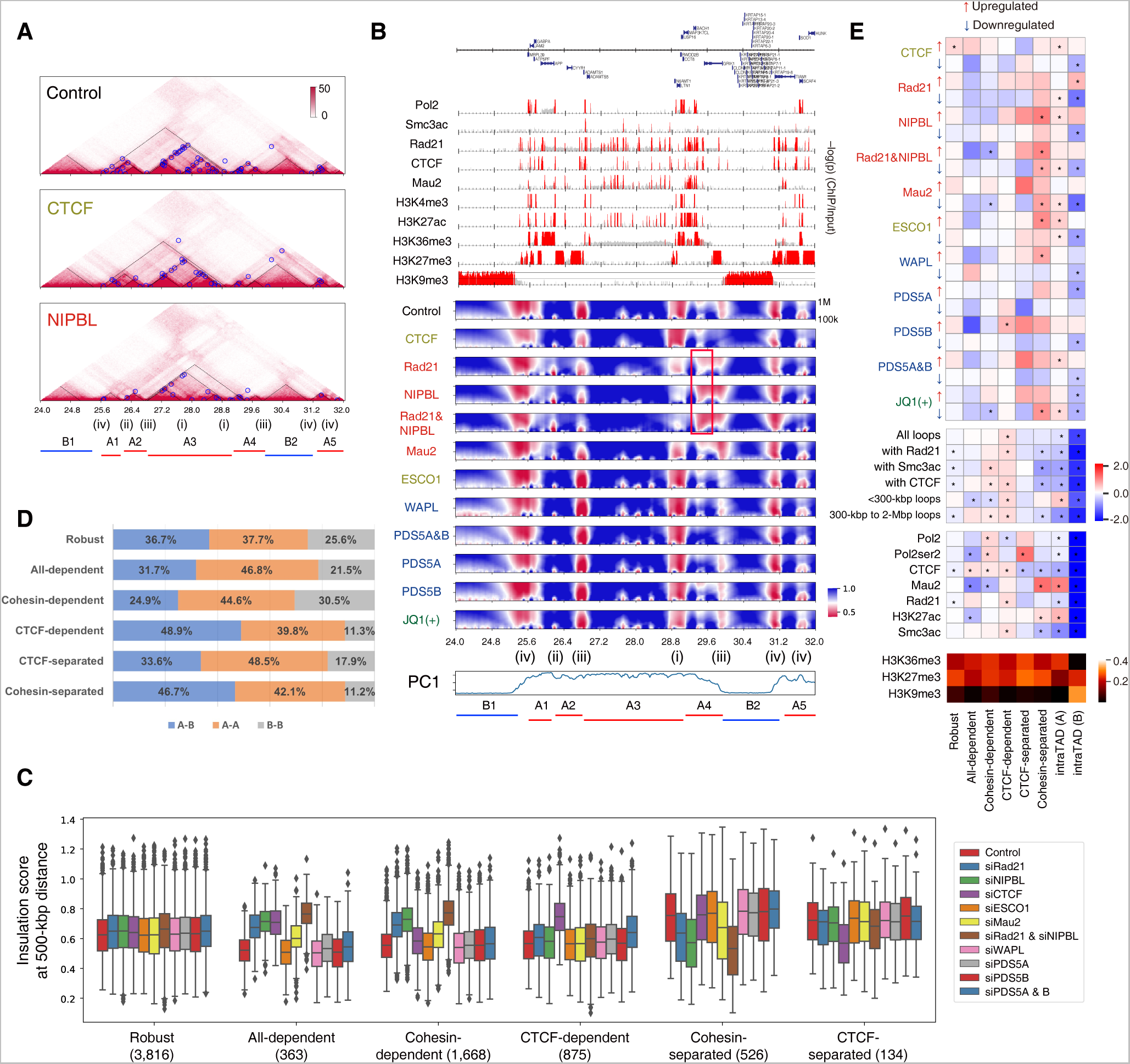
Multi-scale insulation scores reveal the diversity of insulation perturbation at TAD boundaries. **(A-B)** A representative chromosomal region (chromosome 21, 24.0–32.0 Mb). **(A)** Normalized Hi-C matrices. Black dashed lines and blue circles indicate TADs and loops, respectively. **(B)** Top: ChIP-seq distribution (–log_10_(p), 5-kbp bin). Red regions: p < 10^−3^. Middle: Multi-scale insulation scores. Red regions indicate insulated regions (boundaries). The numbers along the bottom (i–iv) indicate four of the six boundary types (see text). The red rectangle indicates the long-range insulation boundaries present after siNIPBL. Bottom: Compartment PC1 and labeled TADs. **(C)** Insulation score distribution for six types of boundaries. The number in parentheses below each boundary type indicates the number of boundaries. **(D)** The proportion of boundaries located between compartments A and B (A-B) or within compartments A (A-A) and B (B-B). **(E)** Relative enrichment of DEGs (top), loops (middle), ChIP-seq peaks (lower middle) and broad histone enrichment (bottom) that overlap the six boundary types and intraTAD regions for compartments A and B against all boundaries. For broad histone marks (bottom), we calculated the fraction of regions covered by the obtained peaks. *p < 0.01, the permutation test (n = 1,000).

We also investigated the overlap of boundaries and ChIP-seq peaks and DEGs (Figures 3E and S6). While cohesin-dependent and CTCF-dependent boundaries were enriched for loops and CTCF peaks, there were few DEGs there. In contrast, cohesin-separated boundaries significantly overlapped with upregulated DEGs after depletion of cohesin and loaders. Upregulated DEGs associated with unloader siRNA were also enriched, although not significantly. At the boundaries, Mau2 was strikingly enriched, but Rad21, CTCF and loops were not. This result suggested that loss of cohesin loading enhances insulation at the Mau2 peak regions (indicating cohesin loading points), which often occur at A-A and A-B boundaries (Figure 3D), resulting in the dysregulation of expression of some genes. As CTCF-separated boundaries also overlapped with DEGs (although not significantly), a gain of boundaries would be more highly correlated with DEGs than a loss of boundaries. Interestingly, DEGs were not enriched at cohesin-dependent boundaries, even though Pol2 was enriched there. Whereas CTCF-independent boundaries were preferred by active genes (Bonev et al. 2017), the loss of such boundaries may not necessarily cause gene expression dysregulation. These observations highlight the necessity of considering the boundary type when investigating the correlation between chromatin folding and gene expression as regulated by cohesin.

### Context-specific depletion effects on interTAD interactions

In addition to the six boundary types (Figure 3C), we also found long-range insulation boundaries that appeared after siNIPBL and siRad21 (500 kbp∼; Figure 3B, red rectangle). The insulation likely reflected the strong depletion of the interaction between an active TAD (A4, enriched by active markers and Pol2) and an inactive compartment B TAD (B2; Figure 4A, black arrows). Although a decreased interaction between active and inactive regions is compatible with finer compartmentalization (Schwarzer et al. 2017), this depletion effect was more region specific and was not symmetric (e.g., there was a milder effect between B2 and A5; Figure 4A). Moreover, we also observed a difference even between siRad21 and siNIPBL on interTAD interactions (e.g., A3-B2; Figure 4A, black rectangles), despite their closely similar effects on TAD and loop structures. We were therefore interested in the variation in perturbations of interTAD interactions among different siRNA targets.

**Figure 4.**
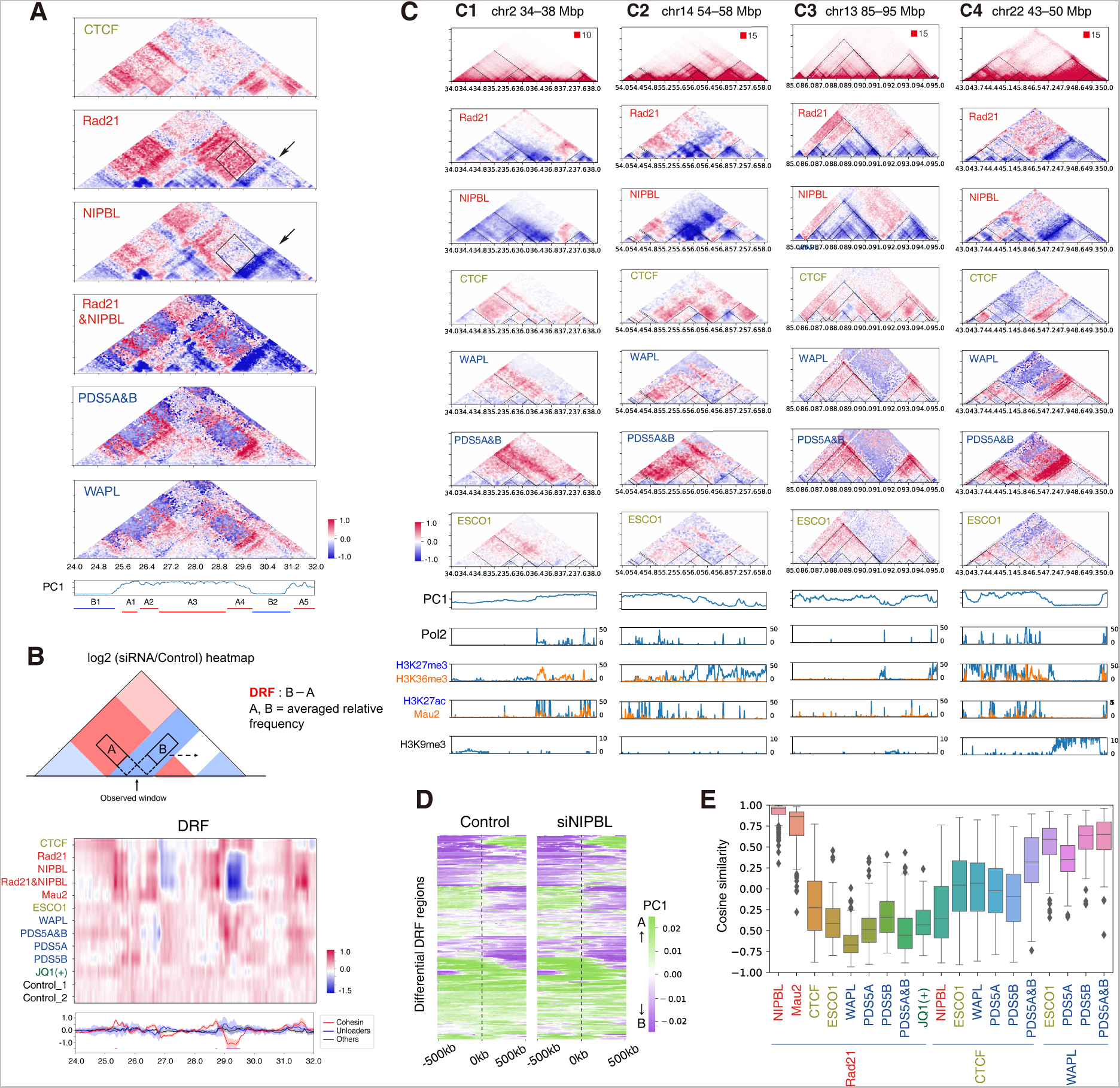
InterTAD-level insulation is increased by cohesin loss. **(A)** Relative enrichment of the interaction frequency (log scale) relative to control. The region and TAD labels (A1–A5, B1 and B2) are as in Figures 3A and 3B. See Figure S7A for all samples. **(B)** Top: Schematic illustration of DRF. Middle: Heatmap of DRF for depletions (row) at the same region with **(A)**. Bottom: Averaged DRF of cohesin and loaders (red), unloaders (blue) and others (black). The shaded regions indicate the 99% confidence interval. The purple bars indicate the identified differential DRF regions. **(C)** Examples of differential DRF regions. Top: Normalized Hi-C matrix (control) and relative interaction frequency after depletions. The dashed black triangles indicate TADs in the control sample. Bottom: Compartment PC1 and ChIP-seq distribution (–log_10_(p), 5-kbp bin). **(D)** Compartment PC1 of the control and siNIPBL sample centered on 241 differential DRF regions. Rows were ordered using hierarchical clustering based on the control sample. **(E)** The cosine similarity distribution of the relative frequency in all 241 differential DRF regions (∼2 Mbp from the center of each region). See Figure S7B for all comparisons.

To identify such a strong effect of depletions on interTAD interactions, we calculated the directional relative frequency (DRF) (Wang and Nakato 2021), which evaluates the directional bias of long-range depletion effects (Figure 4B). We scanned the whole genome and identified 241 regions in which DRF values significantly changed after cohesin or loader depletion (Figure 4C and Table S6). Some of them corresponded to a decrease across broad regions (Figure 4C, C1 and C2), while other regions showed a decreased interaction at one side of TADs (Figure 4C, C3 and C4), reminiscent of the “stripe” structure (Vian et al. 2018). While stripes were reported to be located near super-enhancer regions (Vian et al. 2018), the differential DRF regions in our data were often located at the changing points of compartment PC1 value (Figures 4C and 4D). The strong depletion in interTAD interactions is likely to be distinct from the strengthened compartmentalization because PC1 values around the changing points were not perturbed by siNIPBL (Figure 4D). Moreover, these interactions often increased after siRNA of unloaders; therefore, the effect was inversely correlated. The interaction-increased regions were often larger than the detected TADs in the control (e.g., C1 and C2), suggesting loop extension (Haarhuis et al. 2017; Wutz et al. 2017). Interestingly, siESCO1 showed a smaller but similar effect relative to unloaders in some cases, in which the edge interactions were strengthened (Figure 4C, also supported by the loop length distribution; Figure 1G). We examined this tendency across all 241 regions and confirmed the contrasting depletion effects between cohesin/loaders and unloaders, as well as the positive correlation between ESCO1 and unloaders (except for siPDS5A, Figures 4E and S7B). Considering that ESCO1 facilitates loop stabilization and boundary formation together with CTCF (Wutz et al. 2020), our results suggested that inhibition of cohesin acetylation causes the more frequent pass-through of cohesin at CTCF roadblocks, resulting in loop extension.

### Depletion effects of cohesin and loaders were uncoupled for long-range interactions

We next evaluated the correlation between epigenomic features and depletion effects on long-range interactions using a structured interaction matrix analysis (SIMA) (Lin et al. 2012; Seitan et al. 2013) for interactions at a distance of 500 kbp–5 Mbp. Strikingly, there was a context-dependent difference among depletions (Figures 5A and S8A). Whereas interactions between active markers (H3K4me2, H3K4me3, H3K27ac, Med1 and Pol2) increased after both siRad21 and siNIPBL, interactions between the suppressive marker H3K27me3 and the active markers decreased only after siNIPBL. CTCF depletion increased interactions, especially between promoter marks (Pol2, H3K4me2 and H3K4me3). Again, the depletion of cohesin unloaders showed the opposite tendency relative to that of siNIPBL, whereas SIMA also indicated a difference between PDS5A and B. siPDS5A mainly affected interactions across active markers and H3K27me3, whereas siPDS5B affected cohesin (Rad21 and Smc3ac) and CTCF binding sites, similar to the effect of siWAPL. The effect of siPDS5AB was equivalent to the combined effect of siPDS5A and siPDS5B. siESCO1 affected cohesin, CTCF and enhancer markers.

**Figure 5.**
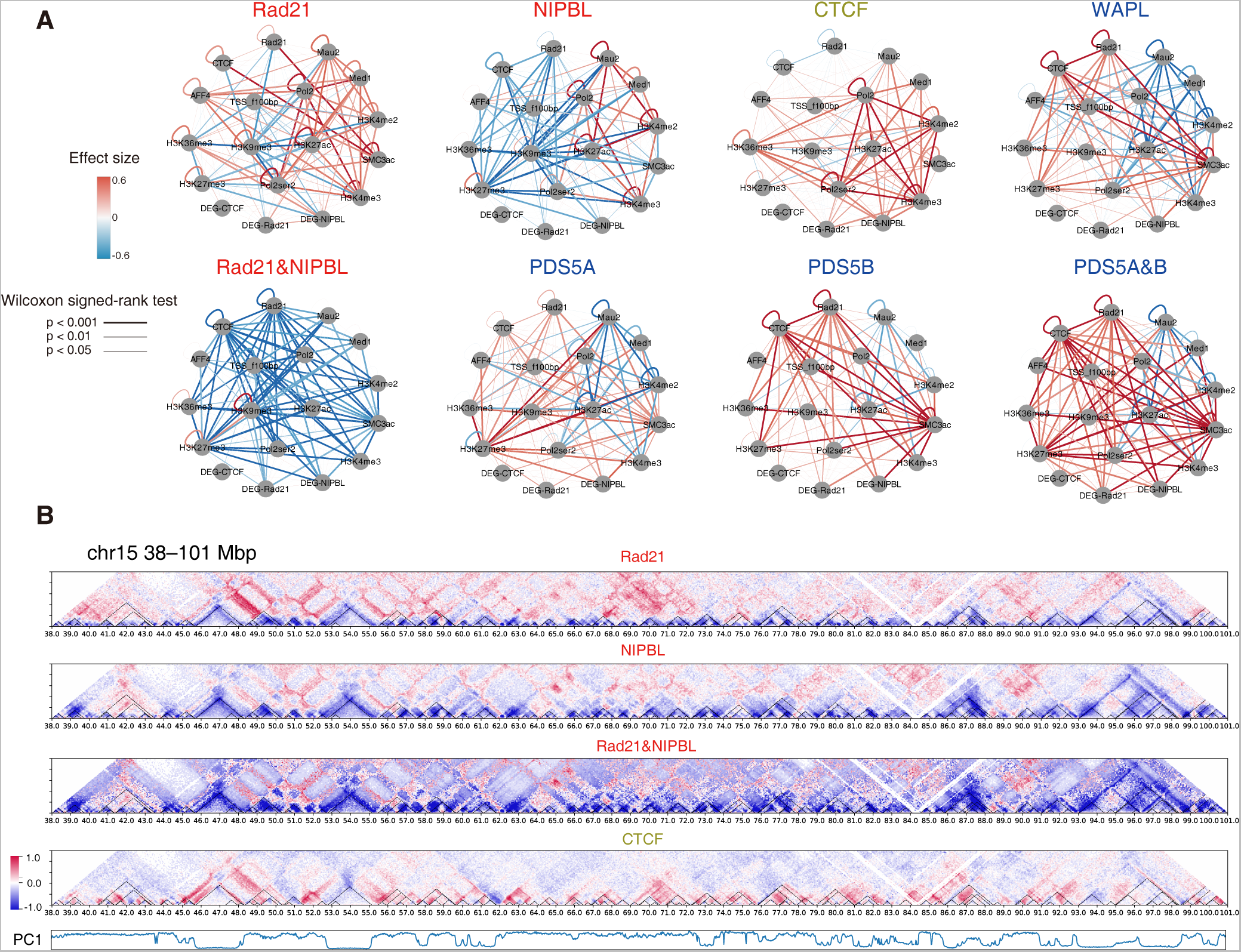
The depletion effect on long-range interactions was different between siRad21 and siNIPBL. **(A)** SIMA analysis after siRNA. See Figure S8A for all samples. **(B)** Relative interaction frequency and compartment PC1 (chromosome 15, 38–101 Mb).

We found that the difference between siRad21 and siNIPBL was mainly derived from global increase of long-range interactions (>2 Mbp) in siRad21, which correspond to interTAD interactions (Figures 5B and S8B). The increase could be involved in strengthened compartmentalization (Rao et al. 2017; Schwarzer et al. 2017), but it cannot explain the difference between siRad21 and siNIPBL. We also found sporadic interactions increased only in siRad21, which might correspond to H3K27ac peaks (Figure 6A). In contrast to our findings in Figure 4, depletion of cohesin unloaders did not show the opposite tendency. Rad21 and NIPBL co-depletion showed a similar effect to siNIPBL alone. This suggested that the amount of cohesin on chromatin has a different effect on long-range interactions relative to the frequency of cohesin loading.

**Figure 6.**
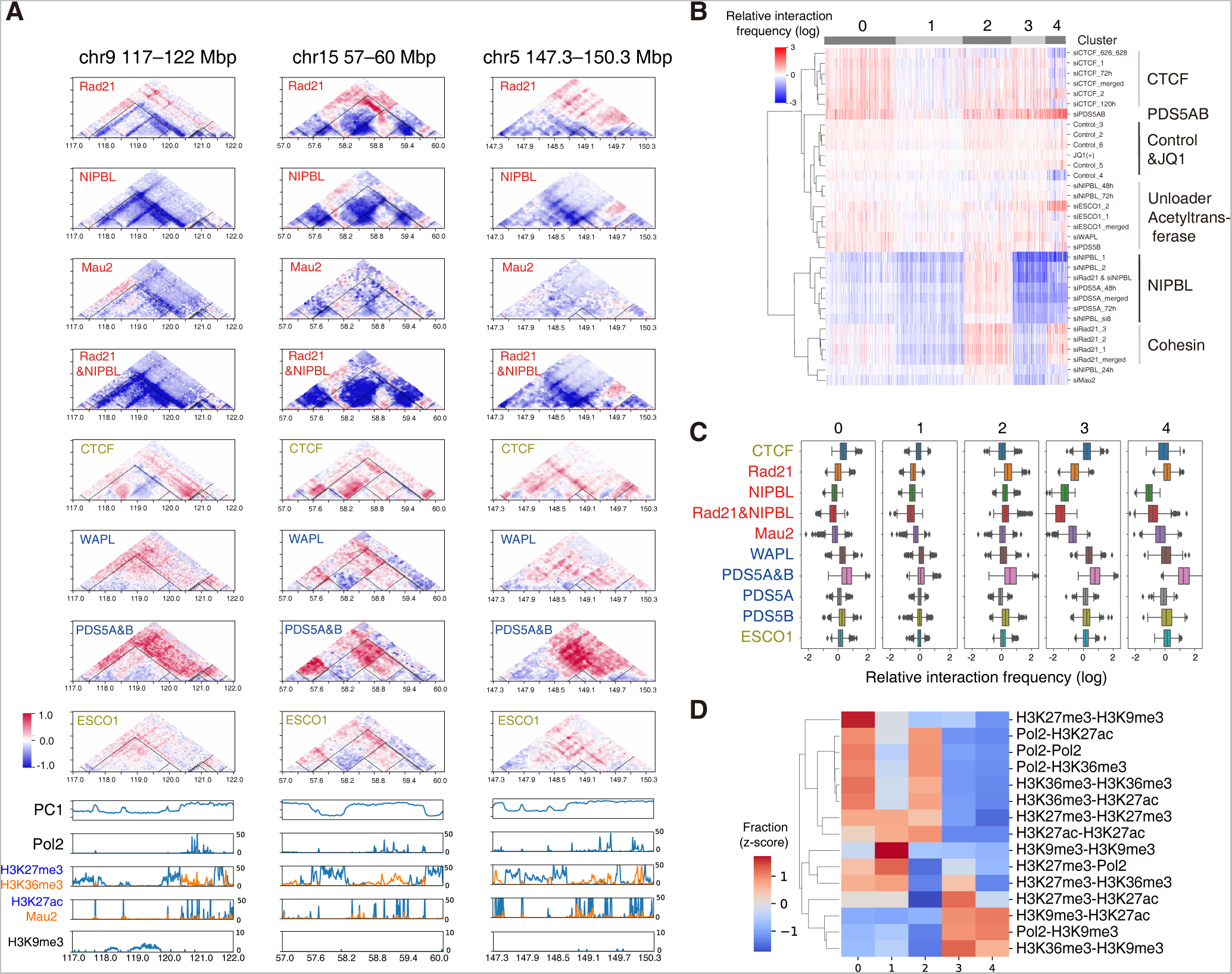
InterTAD interaction variation as compared with epigenomic marks. **(A)** Three example chromosomal regions of increased interactions only in siRad21. Top: Relative frequency and compartment PC1. Bottom: ChIP-seq distribution (–log_10_(p), 5-kbp bin). **(B-D)** K-means clustering (k = 5) of all TAD pairs based on the depletion effect on interactions between them **(B)**, distribution of depletion effects on the clusters **(C)**, and fraction of the epigenomic state of TAD pairs included in the clusters **(D)**.

Finally, we tested whether there is a region-wide depletion effect between TADs and the relationship to the epigenome, as was shown in Figure 4A. For this, we annotated all TADs with epigenomic marks and then classified all TAD pairs based on the relative change in interactions between them (Figures 6B–D). The interactions increased after siRad21 but decreased after siNIPBL in cluster 4. The cluster was enriched for interactions between H3K9me3 and active marks (Figure 6D), which included the difference shown in Figure 4A (A3-B2, black rectangle). The interactions decreased after both siRad21 and siNIPBL in clusters 1 and 3, which included interactions among H3K27me3, H3K9me3 and active markers. This tendency was consistent with the depleted long-range interactions (A4-B2; Figure 4A). In summary, depletion of cohesin and a cohesin loader differently affected the long-range interTAD interactions in an epigenomic-dependent manner.

### Cohesin is broadly distributed but is more important for TAD formation in the active compartment

To directly analyze the depletion effect on the epigenome, we detected the genomic regions significantly changed after siRNA compared with control (Figures 7A and S9A). H3K27ac marks were largely perturbed after NIPBL and Rad21 depletion, whereas H3K36me3 and H3K9me3 were not substantially affected. Though H3K27me3 enrichment also increased after siNIPBL (Figure S9A), we did not observe a spread of it caused by the loss of TAD boundaries. Therefore, the H3K27ac perturbation was likely derived from changes in intraTAD or interTAD interactions.

**Figure 7.**
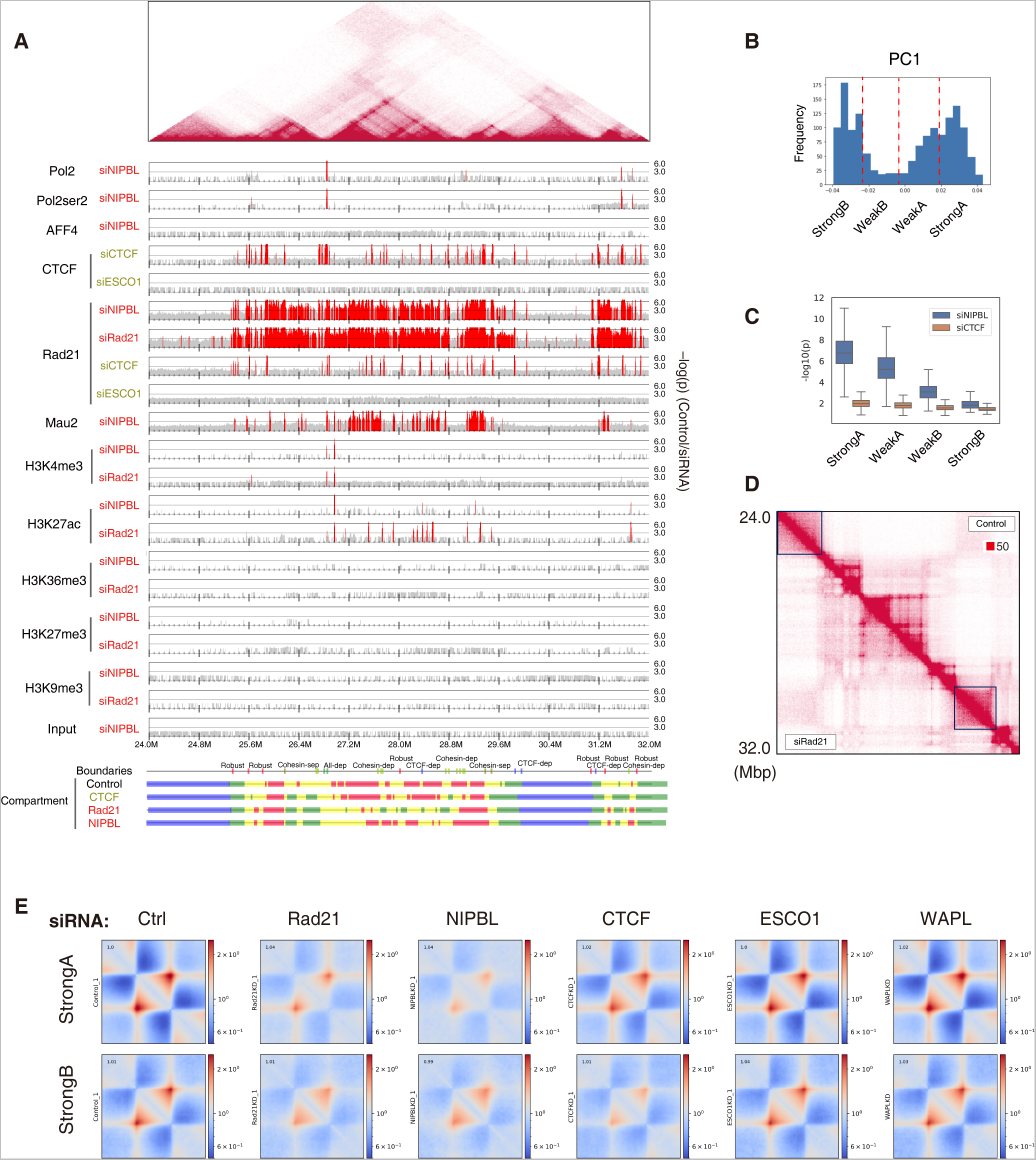
Cohesin was broadly enriched in compartment A but was not enriched in compartment B. **(A)** The depletion effect distribution from ChIP-seq data (–log_10_(p), control/siRNA, 5-kbp bin) on chromosome 21, 24–32 Mbp. The colored bars (bottom) indicate the four compartment types (red, StrongA; yellow, WeakA; green, WeakB; blue, StrongB). **(B)** A graphical representation of the four compartments. **(C)** The –log_10_(p) distribution for the depletion effect of siNIPBL and siCTCF on Rad21 ChIP-seq in the four compartments. **(D)** siRad21 had less of an effect on StrongB TADs (black rectangles). The region is the same with **(A)**. **(E)** Averaged interactions (observed/expected) in StrongA and StrongB TADs for representative samples.

Remarkably, we observed a broad decrease in cohesin after siRad21 and siNIPBL, whereas siCTCF depleted only Rad21 binding at CTCF binding sites (Figure 7A). This indicated that cohesin is located not only in Rad21 peak regions but also in background regions, as assumed in the loop extrusion model (Fudenberg et al. 2016). CTCF acts as an obstacle for cohesin translocation (resulting in sharp cohesin peaks at CTCF sites) but does not control the amount of cohesin on chromatin. This also explains why CTCF depletion decreased loops but had less of an effect on intraTAD interactions (Figures 1D and 1E). However, in compartment B regions, such a substantial depletion of cohesin in the background was not observed (Figure 7A, blue bars). We further investigated this tendency across the genome by dividing compartments A and B into “strong” and “weak” ones based on their PC1 values and confirmed the larger amount of cohesin particularly in “strongA” regions (Figures 7B and 7C). We further investigated the cohesin density using extended ChromHMM (Wang and Nakato 2021) and found that cohesin accumulated to the highest levels at highly active sites (enriched by H3K27ac and H3K4me, Figure S9B). Heterochromatin regions enriched for H3K9me3 in compartment B showed subtle cohesin enrichment. siCTCF did not show such a context-specific tendency (Figures 7C and S9B), and therefore the imbalance in the amount of cohesin between compartments A and B was retained even after loss of CTCF-dependent boundaries, which are often located between A and B (Figure 3D). We also found a milder loss of intraTAD interactions in StrongB than in StrongA, although siCTCF affected both (Figures 7D and 7E). Taken together, our data showed that cohesin also accumulated in non-peak regions, mainly in compartment A, possibly to regulate genes and enhancer activity. In heterochromatic regions, a very small amount of cohesin may be sufficient to maintain TADs, or other systems may maintain TADs such as phase separation (Strom et al. 2017) instead of loop extrusion.

## DISCUSSION

Despite multiple promising models (Wendt et al. 2008; Kagey et al. 2010; Schmidt et al. 2010; Izumi et al. 2015; Fudenberg et al. 2016), the cooperative or distinct roles of cohesin in combination with related factors with respect to chromatin folding and gene expression are not fully understood, especially in a context-specific manner. In this study, we found a variety of TAD boundaries and interTAD interactions that should be considered when investigating the functional and mechanistic relationships of cohesin. Whereas several studies have reported genome clustering based on a single Hi-C data—e.g., a third compartment (Yaffe and Tanay 2011) and six subcompartments (Rao et al. 2014)—our analysis focused on the variation of depletion effects and classified genomic regions using multiple Hi-C data. Although it should be noted that cohesin- and CTCF-independent boundaries may be lost after extreme depletion (e.g., by an auxin-inducible degradation system; (Rao et al. 2017)), our data have delineated the dominant factors for boundaries. The perturbation of long-range interTAD interactions observed in this study cannot be captured by a typical analysis that evaluates only the number/strength of TADs and loops.

Most of the cohesin-related DEGs were related to the direct binding of cohesin around TSSs, which was more CTCF-independent. This result is reminiscent of a report using mouse embryonic fibroblast (MEF) cells (Busslinger et al. 2017), in which cohesin accumulated near TSSs of active genes after CTCF knockout. Some DEGs were also enriched near cohesin-separated boundaries, in which Mau2 specifically accumulated. In contrast, the disruption of TAD boundaries was less correlated with DEGs and histone modifications. We also found DEGs that were similarly dysregulated among siNIPBL and depletion of unloaders, possibly due to dysregulation of cohesin turnover. Because the deficiency of cohesin turnover is one of the causes for CdLS (Deardorff et al. 2012), the DEGs could be candidates for CdLS studies. Whereas the BRD4 mutation also causes a CdLS-like syndrome (Olley et al. 2018), there was little effect of JQ1 on chromatin folding despite the many isolated JQ1-related DEGs. Considering the report that BET inhibition does not disrupt enhancer–promoter contact (Crump et al. 2021), the CdLS phenotype might not involve the perturbation of chromatin structure but could be caused by direct transcription regulation by cohesin, e.g., transcription machinery interacting with Pol2 (Izumi et al. 2015; Busslinger et al. 2017).

For interTAD interactions, although we also found several genes whose changes in expression were consistent with the depletion effect on the interTAD interaction, the effect was often not region-wide (e.g., RUNX1; Figures S9C and S9D). Region-wide gene expression dysregulation may occur with gene clusters: for example, most *PCDH* genes located on chromosome 5 were detected as DEGs and showed cooperative dysregulation (upregulated with unloader depletion, downregulated with loader depletion; Figure S10). Whether these DEGs are the result of TAD disruption or of changes in interTAD interactions, or if their expression is regulated independently via cohesin binding at TSSs or other gene-specific factors, are important questions for future studies.

Despite their closely similar effects on TADs and loops, we also observed a difference in siRad21 and siNIPBL with respect to H3K27me3 and H3K9me3 enrichment. Together with the strengthened compartmentalization under siMau2, the frequency of cohesin loading should be considered separately from the amount of cohesin on chromatin. Meanwhile, the epigenomic state alone cannot fully explain the trend for interTAD interactions, given that the same phenomenon does not occur in all regions that show the same epigenomic pattern.

Compared with the cohesin loaders, the depletion of unloader proteins had different effects on transcriptome and chromatin folding, suggesting their nonredundant roles. WAPL and PDS5B showed a more unloader-like effect, whereas siPDS5A was more related to effects on the epigenome (e.g., Figure 5A). siESCO1 showed a weaker but similar effect relative to that of unloader depletion on chromatin folding, suggesting the involvement of cohesin acetylation in CTCF roadblocks. Considering that DEGs of siESCO1 overlapped to a greater extent with cohesin loader and unloaders than did those associated with siCTCF, the role of cohesin acetylation in gene expression regulation would be mainly related to the cohesin pass-through at CTCF roadblocks in loop extrusion, rather than the formation of stable chromatin loops mediated by acetylated cohesin (Wutz et al. 2020).

We demonstrated that cohesin is broadly distributed within compartment A, not only at peak sites of Rad21 or CTCF localization. Because cohesin should simultaneously function to regulate gene expression and form TADs, a large amount of cohesin could be required to maintain dynamic structures in compartment A. Cohesin did not accumulate substantially in compartment B, in which there was a smaller effect of cohesin depletion. Under the loop-extrusion model, cohesin distribution should have become more uniform after CTCF depletion because of the loss of cohesin stalling at CTCF sites. However, the unequal amounts of cohesin between compartments A and B were retained after siCTCF, whereas we did observe increased interactions between neighboring TADs with the depletion of CTCF-dependent boundaries (e.g., Figure 4A). Therefore, the amount of cohesin on the genome would not be merely derived from loop extrusion but be affected by the genomic context. Moreover, because the interTAD pattern and DEGs of siCTCF were distinct from siRNA of unloaders, the 3D genome segmentation that is retained after siCTCF, and therefore which is likely to be CTCF independent, may be an independent mechanism from the extended TADs observed previously with depletion of unloaders (Haarhuis et al. 2017). How the genomic segmentation within the whole genome is regulated by (extended) loop extrusion and other mechanisms and whether it is conserved among cell types and during cell differentiation remain essential questions for future studies.

## METHODS

### Cell culture and siRNA

We used the siRNA system for depletion, as the auxin-inducible degradation system reduces protein levels even in the absence of auxin, which is not suitable as a control (Wutz et al. 2020). RPE cells (Wendt et al. 2008) were cultured in DMEM (Wako) supplemented with Penicillin-Streptomycin-L-Glutamine Solution (Wako), 10% fetal bovine serum (Biosera) and 20 mM HEPES-KOH (pH 7.4). All siRNA transfections were performed using Lipofectamine RNAiMAX (Thermo Fisher Scientific) in accordance with the manufacturer’s protocol 2 or 3 days before sample preparation, using a final RNA duplex concentration of 50 nM. The siRNA sequences are shown in Supplementary Table S7 and are the same as those described previously (Deardorff et al. 2012; Minamino et al. 2015). For inhibition of BET family proteins, cells were treated with JQ1 for 6 h at a 1 μM final concentration. We labeled JQ1-treated and the corresponding control samples as JQ1(+) and JQ1(-), respectively.

### Antibodies

Antibodies used for ChIP and immunoblotting were as follows. Antibodies against histone H3 lysine-27 acetylation (H3K27ac) (Stasevich et al. 2014), H3 lysine-4 trimethylation (H3K4me3), H3 lysine-9 trimethylation (H3K9me3), H3 lysine-36 trimethylation (H3K36me3) and Pol2ser2 were provided by Dr. Kimura (Tokyo Institute of Technology, Tokyo, Japan). We also used antibodies against Rad21, Smc3ac and ESCO1, which were described previously (Minamino et al. 2015). Antibodies against NIPBL (A301-779A, BETHYL), Mau2 (ab46906, abcam), CTCF (07-729, Merck), BRD4 (A301-985A50, BETHYL), AFF4 (A302-538A, BETHYL), Pol2 (14958, Cell signaling technology) and H3 lysine-27 trimethylation (H3K27me3, ab192985, abcam) were used for ChIP. Antibodies against NIPBL (sc-374625, Santa Cruz Biotechnology), a-tubulin (T6074, Merck), WAPL (16370-1-AP, Proteintech), PDS5A (A300-088A, BETHYL), PDS5B (ab70299, abcam) and CTCF (3417, Cell signaling technology) were used for immunoblotting. The mouse monoclonal antibody against Mau2 was generated using a synthetic peptide corresponding to residues 596-613 (PVQFQAQNGPNTSLASLL) of human Mau2 and used for immunoblotting. Antibody dilutions in immunoblotting were 1:500 (NIPBL and ESCO1) and 1:1,000 (other antibodies).

### Protein analysis

Cells were lysed with lysis buffer (20 mM HEPES-KOH, pH 7.5; 100 mM NaCl; 10 mM KCl; 10% glycerol; 340 mM sucrose; 1.5 mM MgCl_2_; 10 mM sodium butyrate; 0.25% Triton X-100; 1 mM dithiothreitol; 1× cOmplete proteinase inhibitor cocktail) as described (Deardorff et al. 2012). The resulting lysate was mixed with SDS-PAGE sample buffer (50 mM Tris-HCl, pH 6.8; 2% SDS; 0.05 % BPB; 7% glycerol; 5% 2-mercaptoethanol) and boiled for 5 min. The proteins were analyzed by Mini-PROTEAN Tetra Vertical Electrophoresis Cell (Bio-Rad) in accordance with the manufacturer’s protocol.

### *In situ* Hi-C

We used the *in situ* Hi-C protocol as described in Rao *et al*. (Rao et al. 2014). In brief, ∼3 × 10^6^ RPE cells were crosslinked with 1% formaldehyde for 10 min at room temperature, followed by an additional 5 min with 200 mM glycine in phosphate-buffered saline (PBS). Fixed cells were permeabilized in Hi-C lysis buffer (10 mM Tris-HCl, pH 8.0; 10 mM NaCl; 0.2% Igepal CA630; 1× protease inhibitor cocktail [Sigma]) on ice. The cells were treated with 100 U of MboI (New England Biolabs) for chromatin digestion, and the ends of digested fragments were labeled with biotinylated nucleotides followed by ligation. After DNA reverse crosslinking and purification, ligated DNA was sheared to a size of 300–500 bp using a Covaris S2 focused-ultrasonicator (settings: Duty Cycle, 10%; Intensity, 4; Cycles per Burst, 200; Duration, 55 sec). The ligated junctions were then pulled down with Dynabeads MyOne Streptavidin T1 beads (Thermo Fisher Scientific). The pulled-down DNA was end-repaired, ligated to sequencing adaptors, amplified on beads and purified using Nextera Mate Pair Sample Preparation Kit (Illumina) and Agencourt AMPure XP (Beckman Coulter). DNA was then sequenced to generate paired-end 150-bp reads using the Illumina HiSeq-2500 or X Ten system.

### Hi-C data processing with Juicer

Sequenced reads were processed using Juicer version 1.5.7 and Juicer tools version 1.9.9 (Durand et al. 2016), and the definition of TADs and loops follows Rao *et al*. (Rao et al. 2014). The detailed steps are as follows: Sequenced paired-end reads were mapped by BWA version 0.7.17 (Li and Durbin 2009). We generated contact map files with square root vanilla coverage (VC_SQRT) normalization. We used 25-kbp resolution maps unless otherwise described. We called TADs using Arrowhead. Because the obtained TADs can be nested, we also generated a list of non-overlapping TADs by segmenting the genome based on all TAD boundaries. TAD boundaries were defined as edges for all annotated TADs. Loops were called at 5-kbp, 10-kbp and 25-kbp resolution by HiCCUPS. To obtain peak-overlapping loops (Figure 2C), we used BEDTools v2.28.0 (https://bedtools.readthedocs.io/en/latest/) and extracted loops for which both anchor sites overlapped with the peaks. CTCF motif analysis was implemented using MotifFinder. High-resolution data that combined all replicates were generated by *mega.sh* script provided by Juicer. Eigenvector (PC1) values for compartment analysis were generated with the *eigenvector* command in Juicertools. For allele-specific Hi-C analysis of chromosome X, we obtained single-nucleotide polymorphism data for RPE cells from Darrow *et al. (Darrow et al. 2016)*, which was then converted to genome build hg38 by the liftOver tool (https://genome-store.ucsc.edu/). We modified the *diploid.sh* script provided by Juicer and made interaction map files for active and inactive chromosome X.

The samples and mapping statistics are summarized in Tables S1. Since we generated six replicates as control samples, we merged them into a single high-resolution Hi-C data and used it to obtain reference data for the TADs, loops and compartment data. For the comparative analysis, we normalized Hi-C matrices based on the number of mapped reads on each chromosome. Therefore, the tendency for increases and decreases is relative; that is, increased long-range interactions might be compensated for by increased short-range interactions (Nora et al. 2020). siRad21 and siNIPBL were most affected by this fact, because almost all TADs and loops were depleted after these treatments. Therefore, in our analysis, we focused on the variation of depletion effects across samples to capture the context-specific tendency, rather than translating the biological meaning of increased/decreased interaction itself.

### Hi-C data processing with other tools

To evaluate the quality and reproducibility of our Hi-C data, we used 3DChromatin_ReplicateQC (Yardimci et al. 2019), which internally implements QuASAR (Sauria and Taylor 2017) and HiCRep (Yang et al. 2017). Because of the large computational complexity involved, we used only chromosomes 21 and 22 with a 50-kbp bin for the quality evaluation. We confirmed that all of our Hi-C data had sufficient quality (QuASAR-QC scores > 0.05, Table S1). HiCRep was used to evaluate the overall similarity of the depletion effects among our Hi-C samples by calculating a stratum-adjusted correlation coefficient that captures the similarity of chromatin features including TADs and loops. We used Cooler (Abdennur and Mirny 2020) and cooltools (https://cooltools.readthedocs.io/) for APA plots, averaged TAD plots and saddle plots. The compartment strength was defined as (AA + BB) –2(AB), where AA, BB and AB indicate the interaction frequency between compartments A and A, compartments B and B and compartments A and B, respectively, of the saddle plot. The visualization of Hi-C matrices with ChIP-seq distributions were visualized using Python.

### Structured interaction matrix analysis

To explore interactions between specific chromatin features (e.g., ChIP-seq peaks), we used structured interaction matrix analysis (SIMA) (Lin et al. 2012) implemented in HOMER (http://homer.ucsd.edu/homer/). SIMA assembles information for multiple occurrences of each feature, providing an overview of Hi-C interactions associated with a genomic feature between each pair of specified domains. In this study, the genomic features included the ChIP-seq peak list (AFF4, CTCF, H3K27ac, H3K27me3, H3K36me3, H3K4me3, H3K4me2, Smc3ac, Mau2, Med1, Pol2, Pol2ser2 and Rad21), differentially expressed genes (DEGs) (siCTCF, siNIPBL, siRad21) and transcriptional start sites (TSSs). Domains of interest were defined as TAD lists, and the distances between two TADs were specified to be <5 Mbp, <2 Mbp or 2–<5 Mbp with ‘-max -min’ parameters. By making comparisons with the background model, we obtained an enrichment score, representing the degree to which a genomic feature pair was enriched in the Hi-C interactions between two TADs. To compare differences in enrichment scores between cohesin-knockdown and control Hi-C samples, we used the paired Wilcoxon signed-rank test to calculate the p-value and effect size for each genomic feature pair, as described (Seitan et al. 2013). We used Cytoscape (Shannon et al. 2003) to visualize the results.

### Multi-scale insulation score

A multi-scale insulation score was generated as described (Crane et al. 2015). In brief, the insulation score was calculated at a resolution of 25 kbp as the log-scaled relative contact frequency across pairs of genomic loci located around the genomic positions from 100 kbp to 1 Mbp. The 500-kbp distance was used in the insulation score analysis. For the classification of insulation boundaries into six types, we used the following criteria based on this 500-kbp insulation score:

1. if(siNIPBL – control) > T_ins_ and if(siCTCF – control) > T_ins_ : “all-dependent”
2. else if(siNIPBL – control) > T_ins_ or if(siRad21 – control) > T_ins_ : “cohesin-dependent”
3. else if(siCTCF – control) > T_ins_ : “CTCF-dependent”
4. else if(control – siCTCF) > T_ins_ : “CTCF-separated”
5. else if(control – siNIPBL) > T_ins_ or if(control – siRad21) > T_ins_ : “cohesin-separated”
6. else if(siNIPBL – control) < T_ins_ and if(siCTCF – control) < T_ins_ and if(siCTCF – control) < T_ins_ : “robust”

where we set T_ins_, the threshold of insulation score, as 0.13. We excluded siMau2 as a criterion because it had a smaller effect than siNIPBL and siRad21 on the insulation score. We excluded chromosomes X and Y and the mitochondrial genome from this boundary analysis. The obtained six types of boundaries are summarized in Table S5.

### Directional relative frequency (DRF)

DRF measures the bias in the relative interaction frequency (*T* = *log* (*C_siRNA_*) – *log* (*C_control_*)) between regions up- and downstream of each genomic region, where *C* is a normalized contact matrix. Therefore, the DRF can be calculated by

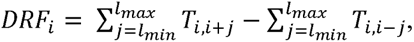

where *l_min_* and *l_max_* indicate the range of interaction. In this study, we set *l_min_* = 500 *kbp* and *l_max_* = 2 *Mbp*.

To obtain differential DRF regions, we classified Hi-C samples into “cohesin and loaders,” “cohesin unloaders” and “others (including control)” and calculated the averaged DRF values and a 99% confidence interval (CI). Then we identified the regions that satisfied the following criteria: the 99% CI ranges of “cohesin and loaders” and “others” did not overlap, and the averaged DRF value of “cohesin and loaders” was > T_DRF_ or < -T_DRF_, where T_DRF_ refers to the threshold of DRF. We set T_DRF_ = 0.7 in this study. The obtained differential DRF regions are summarized in Table S6.

### RNA-seq

Total RNA was isolated using Trizol (Thermo Fisher Scientific) and a Nucleospin RNA kit (Macherey-Nagel). rRNA was removed with the Ribo-Zero Gold rRNA Removal Kit (Illumina), followed by sequencing library preparation with the NEBNext Ultra Directional RNA Library Prep Kit for Illumina (New England Biolabs). Single-end 65-bp reads were sequenced by Illumina HiSeq-2500 system. Sequenced reads were mapped to the human reference sequence (GRCh38) by STAR version 2.7.3a (Dobin et al. 2013) with the following options “SortedByCoordinate --quantMode TranscriptomeSAM --outSAMattributes All”. The samples and mapping statistics are summarized in Tables S2. The gene expression levels were estimated by RSEM version 1.3.1 (Li and Dewey 2011) with the option “--estimate-rspd --strandedness reverse”. We used DESeq2 (Love et al. 2014) to identify DEGs (protein-coding genes, false discovery rate [FDR] < 0.01). We focused on protein-coding genes to avoid the effects of repetitive non-coding RNAs.

To mitigate the indirect effect and the technical variances, we generated the list of DEGs by merging the top-ranked 1,000 DEGs from each pairwise comparison between each siRNA and the controls. We used clusterProfiler (Yu et al. 2012) for the GO enrichment analysis.

### Spike-in ChIP-seq

Spike-in ChIP-seq enables us to explore the absolute-level difference in read enrichment among samples (Bonhoure et al. 2014; Nakato and Shirahige 2017). Chromatin preparation for ChIP was performed as described (Izumi et al. 2015). In brief, ∼8 × 10^6^ RPE cells were crosslinked with 1% formaldehyde for 10 min at room temperature, followed by an additional 5 min with glycine in PBS added at a final concentration of 125 mM. Fixed cells were lysed in LB1 (50 mM HEPES-KOH, pH 7.4; 140 mM NaCl; 1 mM EDTA; 10% glycerol; 0.5% NP-40; 0.25% Triton X-100; 10 mM dithiothreitol; 1 mM PMSF) on ice. The lysate was washed sequentially with LB2 (20 mM Tris-HCl, pH 8.0; 200 mM NaCl; 1 mM EDTA; 0.5 mM EGTA; 1 mM PMSF) and LB3 (20 mM Tris-HCl, pH 7.5; 150 mM NaCl; 1 mM EDTA; 0.5 mM EGTA; 1% Triton X-100; 0.1% sodium deoxycholate; 0.1% SDS; 1× cOmplete protease inhibitor cocktail [Roche]) on ice. The lysate was resuspended in LB3 and sonicated using Branson Sonifier 250D (Branson) for chromatin shearing (12 sec with amplitude setting at 17% of the maximum amplitude, six times). In addition, lysate containing fragmented chromatin was also prepared from ∼2 × 10^6^ mouse cells (C2C12) with the same procedures. Human cell lysate and mouse cell lysate (as a spike-in internal control) were combined (∼4:1 ratio) and incubated with protein A or G Dynabeads (Thermo Fisher Scientific) conjugated with the relevant antibodies for 14 h at 4°C. The beads were then washed five times with cold RIPA wash buffer (50 mM HEPES-KOH, pH 7.4; 500 mM LiCl; 1 mM EDTA; 0.5% sodium deoxycholate; 1% NP-40) and once with cold TE50 (50 mM Tris-HCl, pH 8.0; 10 mM EDTA). Material captured on the beads was eluted with TE50 containing 1% SDS. The eluted material and input were incubated for 6 h at 65°C to reverse crosslinks and were treated with 100 ng RNaseA (Roche) for 1 h at 50°C, followed by treatment with 100 ng Proteinase K (Merck) overnight at 50°C. The input and ChIP DNA were then purified with a PCR purification kit (Qiagen). DNA from the ChIP and input fractions was end-repaired, ligated to sequencing adaptors, amplified and size-selected using NEBNext Ultra II DNA Library Prep Kit for Illumina (New England Biolabs) and Agencourt AMPure XP (Beckman Coulter). DNA was then sequenced to generate single-end 65-bp reads using the Illumina HiSeq-2500 and NextSeq 2000 systems.

Reads were aligned to the human genome build hg38 and mouse genome build mm10 using Bowtie2 version 2.4.1 (Langmead and Salzberg 2012) with default parameters. Quality assessment was performed with SSP version 1.2.2 (Nakato and Shirahige 2018) and DROMPAplus version 1.12.1 (Nakato and Sakata 2020). Spike-in read normalization, peak calling and visualization were performed with DROMPAplus. The mapping statistics, quality values and the scaling factors for spike-in normalization are summarized in Table S3. The default parameter set was used for peak calling (100-bp bin, --pthre_internal 5, --pthre_enrich 4) except for H3K9me3 (--pthre_internal 1 -- pthre_enrich 2) due to the lower signal-to-noise ratio. For read visualization (Figures 2–4 and 6), we displayed –log_10_(p) scores of ChIP/input enrichment (--showpenrich 1 option), which is recommended for distinguishing the signal from the noise (Roadmap Epigenomics et al. 2015). For Figure 7, to look for significant changes in ChIP-seq data, we similarly compared ChIP (control) and ChIP (siRNA) and visualized –log_10_(p).

### Permutation test for overlapping analysis

To compare the overlapping ratio of TSSs of DEGs and ChIP-seq peaks, Hi-C loops and insulation boundaries, we implemented the permutation test (n = 1,000) that compared the relative overlap frequency against the background distribution. As a background, we used all DEGs obtained by DESeq2 (11,345 genes, FDR < 0.01) for DEGs, and all boundaries (7,421) for the six types of boundaries. We randomly picked up the same number of genes or boundaries from the background in each permutation and generated the frequency distribution. For the boundary analysis (Figure 3E), we counted DEGs and the peaks that overlapped within 50 kbp of them.

### Correlation of interactions with epigenomes

For interTAD interaction comparisons (Figures 6B–D), we extracted all TAD regions with widths of >100 kbp and annotated them using the epigenomic marks (H3K36me3, H3K27me3, H3K9me3 and Pol2) with the following criterion: whether the marks covered > 40.0% of the TAD length. To avoid a low read coverage at long-range distances and the technical effect derived from the different resolution of Hi-C matrices, we used the log-fold change *log*_2_(*N_siRNA_/N_control_*), where *N_siRNA_* and *N_control_* indicate the total number of fragments mapped within the interTAD regions between a TAD pair (≤2 Mbp in distance) annotated with the epigenomic marks. We calculated the score for all TAD pairs and applied k-means clustering (k = 5). Then we calculated the z-score–normalized fraction of epigenomic status for the TAD pairs included in each cluster to estimate the epigenomic-dependent depletion effect of interTAD interactions.

### Extended ChromHMM

Our previous study showed that several 1D metrics for Hi-C data are effective for annotating chromatin states in detail (Wang and Nakato 2021). In this study, we added CTCF, boundary and compartment information in addition to five core histone marks (H3K4me3, K3K27ac, H3K27me3, H3K36me3 and H3K9me3) to ChromHMM (Ernst and Kellis 2012) and annotated 15 chromatin states.

## DATA ACCESS

The raw sequencing data and processed files of Hi-C, RNA-seq and ChIP-seq data from this study have been submitted to the Gene Expression Omnibus (GEO) under the accession number GSE196450. Custom code used for the principal analysis is available at Docker Hub (https://hub.docker.com/r/rnakato/juicer) and at GitHub (https://github.com/rnakato/RPE_Hi-C_Analysis).

## ACKNOWLEDGEMENTS

We thank all members of the Nakato and Shirahige Laboratories for discussions and comments on the manuscript. This work was supported by a Grant-in-Aid for Scientific Research (17H06331 to R.N. and 15H05970, 20H05686 and 20H05940 to K.S), the Japan Agency for Medical Research and Development under grant number JP22gm6310012h0003 and the Japan Science and Technology Agency under grant number JPMJCR18S5.

## AUTHOR CONTRIBUTIONS

R.N. conceived this project and wrote the manuscript. R.N., J.W., L.A.E.N. and G.M.O. implemented the computational analysis. T.S. prepared Hi-C, ChIP-seq and RNA-seq samples. M.B. prepared ChIP-seq and RNA-seq samples. K.S. supervised the sample preparation and sequencing and suggested ways to improve the analysis and the manuscript.

