## Supplemental Figures for "Context-dependent 3D genome regulation by cohesin and related factors"

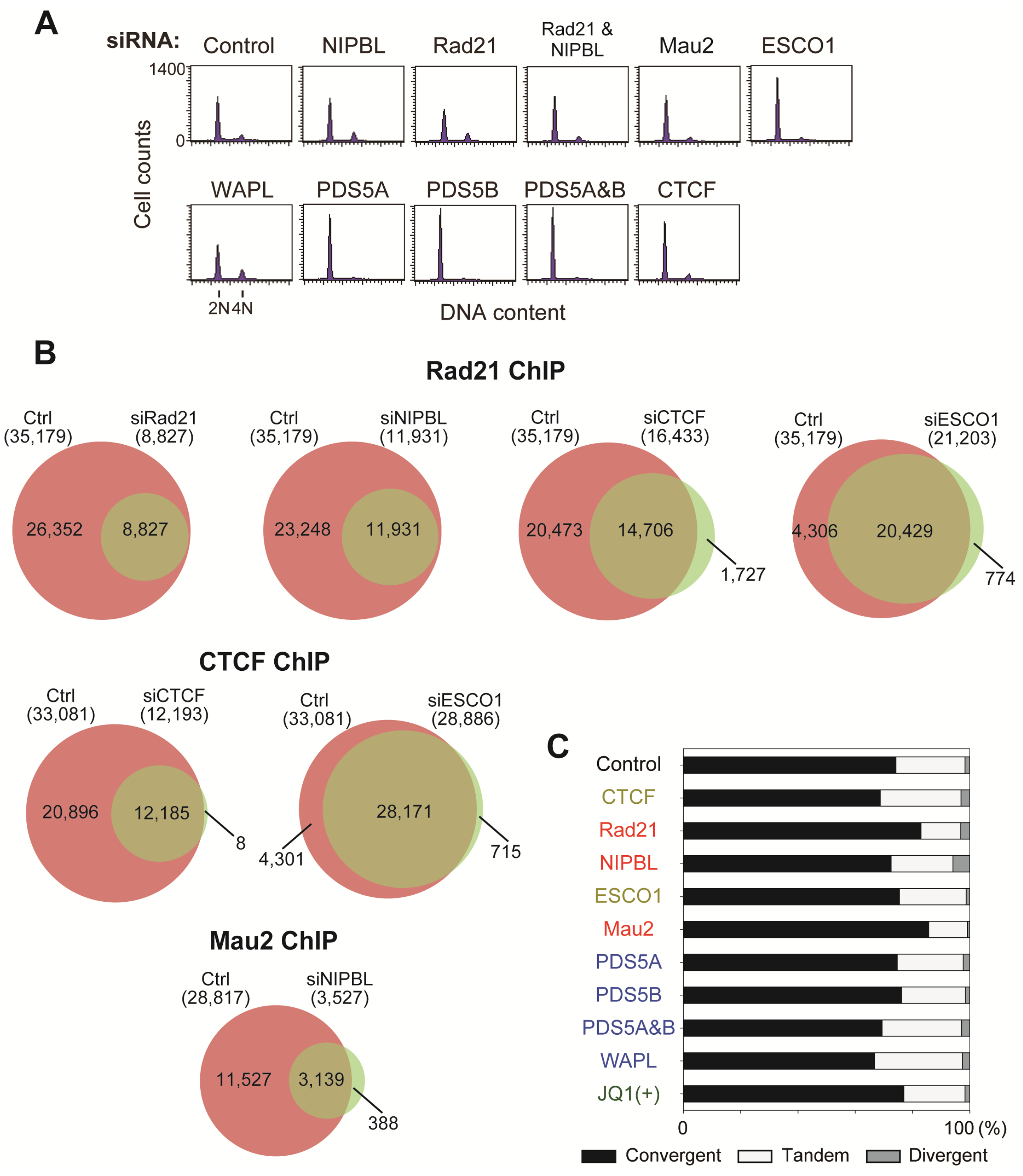


**Figure S1. Depletion effects on ChIP-seq and Hi-C. (A)** Fluorescence-activated cell sorting plots for all depletions. Representative samples are shown. **(B)** Peak overlap between control and siRNA samples. **(C)** The proportion of convergent, tandem and divergent CTCF motif orientations for loops that contained CTCF peaks at the anchor sites among control and siRNA samples.

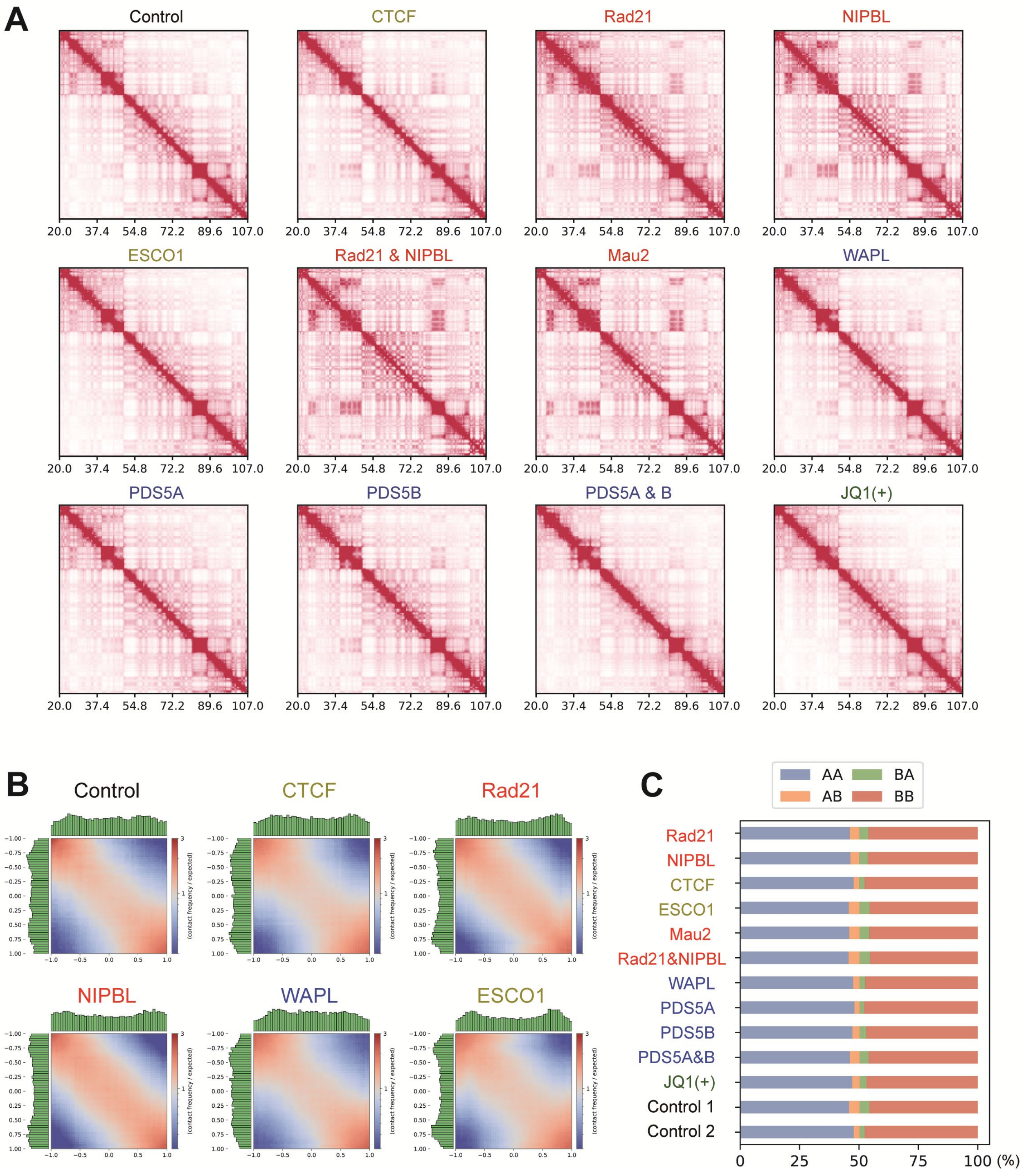


**Figure S2. Compartment strength and switching. (A)** Hi-C maps of a representative chromosomal region (chromosome 14, 20.0–107.0 Mb). Depletion of cohesin and loaders resulted in increases in long-range interactions within compartments A and B, visible here as a plaid (or checkerboard) pattern. **(B)** Saddle plots evaluated compartmentalization strength. Averaged interaction frequencies between pairs of loci sorted by their compartment PC1 values (green histogram). **(C)** The fraction of compartment switching. A, compartment A; B, compartment B. Two control replicates are shown as negative control.

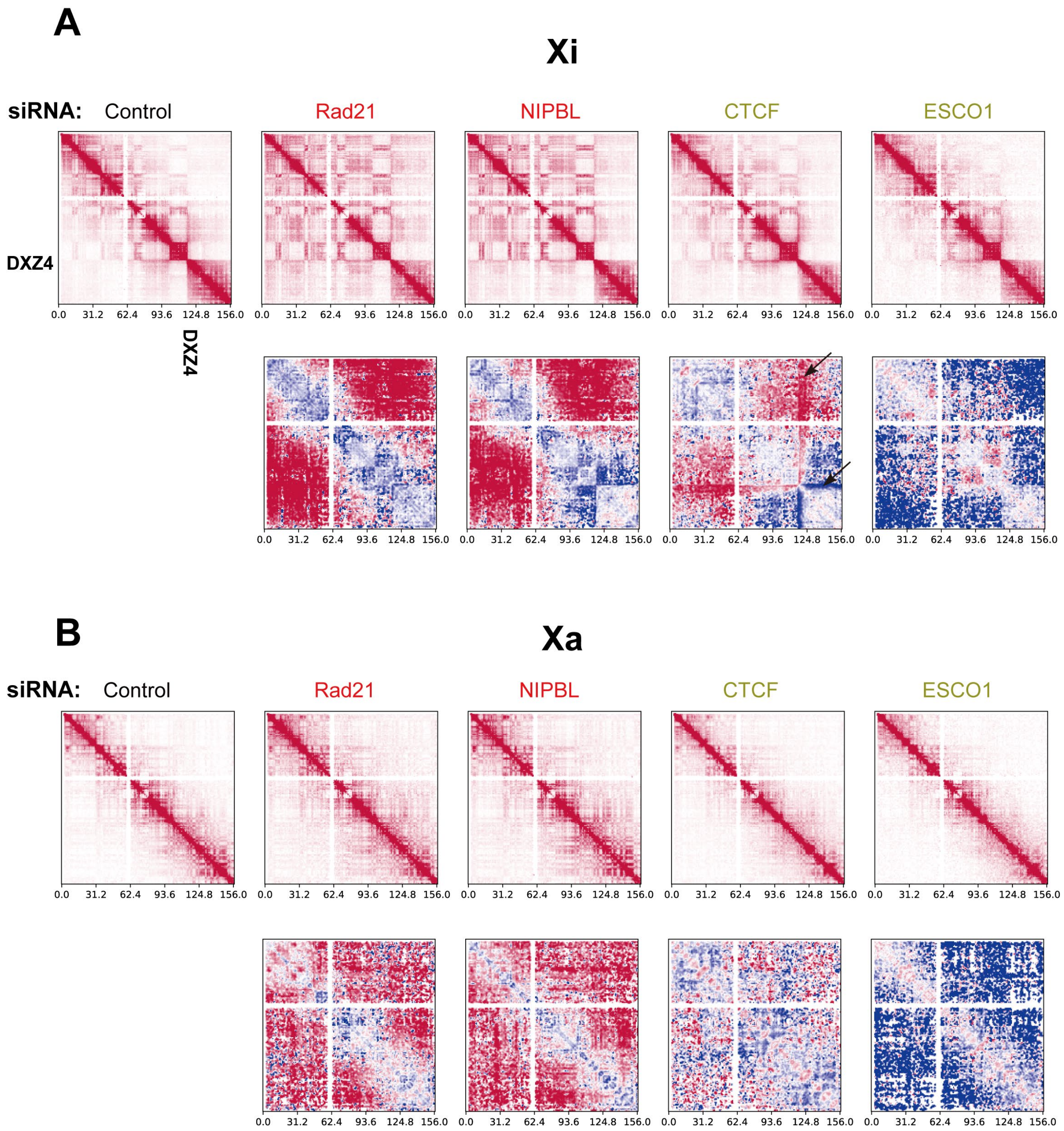


**Figure S3. Depletion effect on active and inactive chromosome X. (A, B)** Allele-specific Hi-C analysis of **(A)** inactive chromosome X (Xi) and **(B)** active chromosome X (Xa). The microsatellite repeat DXZ4 indicates the boundary of the two megadomains. Top: Normalized heatmaps. Bottom: Relative enrichment of the interaction frequency (log scale) relative to control. In Xi **(A)**, the interaction within the smaller megadomain (lower right of the heatmaps) was less affected in all samples.

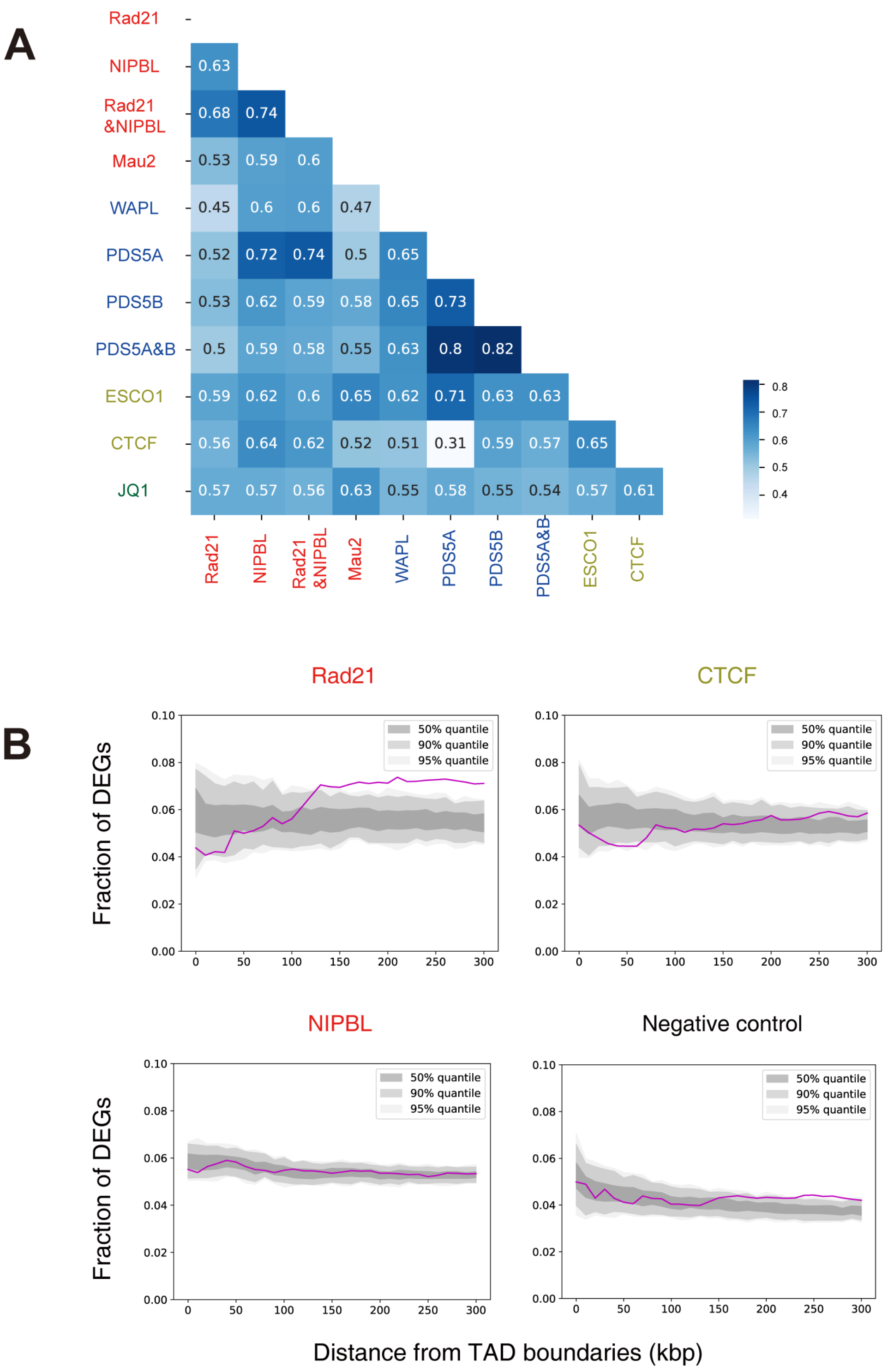


**Figure S4. Comparative analysis of siRNA effects on the transcriptome. (A)** Correlation heatmap based on Simpson index (overlap of all DEGs, FDR < 0.01) for all sample pairs. **(B)** The proximity of DEGs to disrupted TAD boundaries for siRad2, siCTCF, siNIPBL and the negative control (DEGs from siMau2 against disrupted boundaries from siCTCF). Purple lines indicate fraction of DEGs at varying distances from disrupted TAD boundaries. Gray ribbons indicate the 50, 90 and 95% percentiles from random boundaries.

A

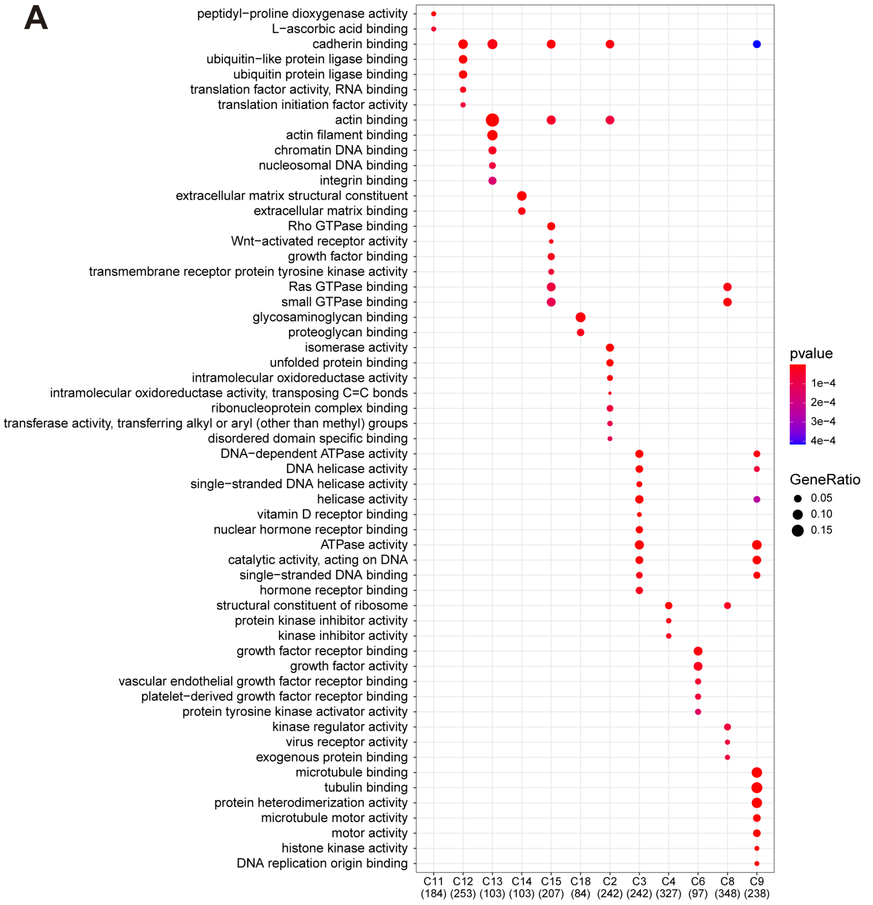
peptidyl−proline dioxygenase activity
L−ascorbic acid binding
cadherin binding
ubiquitin−like protein ligase binding
ubiquitin protein ligase binding
translation factor activity, RNA binding
translation initiation factor activity
actin binding
actin filament binding
chromatin DNA binding
nucleosomal DNA binding
integrin binding
extracellular matrix structural constituent
extracellular matrix binding
Rho GTPase binding
Wnt−activated receptor activity
growth factor binding
transmembrane receptor protein tyrosine kinase activity
Ras GTPase binding
small GTPase binding
glycosaminoglycan binding
proteoglycan binding
isomerase activity
unfolded protein binding
intramolecular oxidoreductase activity
intramolecular oxidoreductase activity, transposing C=C bonds
ribonucleoprotein complex binding
transferase activity, transferring alkyl or aryl (other than methyl) groups
disordered domain specific binding
DNA−dependent ATPase activity
DNA helicase activity
single−stranded DNA helicase activity
helicase activity
vitamin D receptor binding
nuclear hormone receptor binding
ATPase activity
catalytic activity, acting on DNA
single−stranded DNA binding
hormone receptor binding
structural constituent of ribosome
protein kinase inhibitor activity
kinase inhibitor activity
growth factor receptor binding
growth factor activity
vascular endothelial growth factor receptor binding
platelet−derived growth factor receptor binding
protein tyrosine kinase activator activity
kinase regulator activity
virus receptor activity
exogenous protein binding
microtubule binding
tubulin binding
protein heterodimerization activity
microtubule motor activity
motor activity
histone kinase activity
DNA replication origin binding
C11 (184)
C12 (253)
C13 (103)
C14 (103)
C15 (207)
C18 (84)
C2 (242)
C3 (242)
C4 (327)
C6 (97)
C8 (348)
C9 (238)
pvalue
1e−4
2e−4
3e−4
4e−4
GeneRatio
0.05
0.10
0.15

B

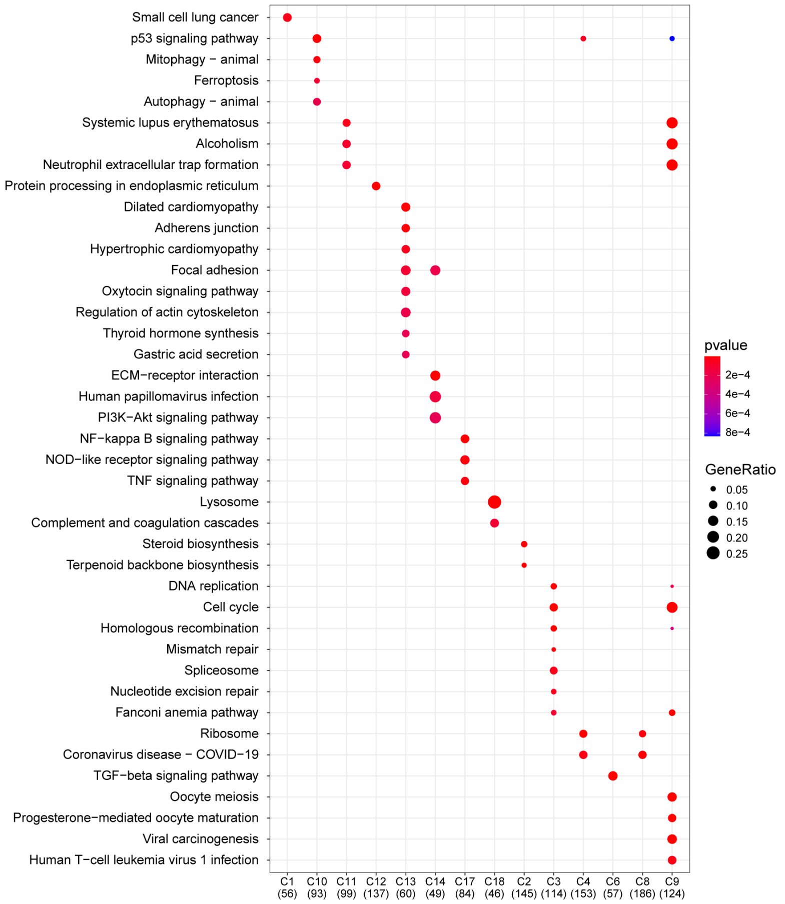
Small cell lung cancer
p53 signaling pathway
Mitophagy − animal
Ferroptosis
Autophagy − animal
Systemic lupus erythematosus
Alcoholism
Neutrophil extracellular trap formation
Protein processing in endoplasmic reticulum
Dilated cardiomyopathy
Adherens junction
Hypertrophic cardiomyopathy
Focal adhesion
Oxytocin signaling pathway
Regulation of actin cytoskeleton
Thyroid hormone synthesis
Gastric acid secretion
ECM−receptor interaction
Human papillomavirus infection
PI3K−Akt signaling pathway
NF−kappa B signaling pathway
NOD−like receptor signaling pathway
TNF signaling pathway
Lysosome
Complement and coagulation cascades
Steroid biosynthesis
Terpenoid backbone biosynthesis
DNA replication
Cell cycle
Homologous recombination
Mismatch repair
Spliceosome
Nucleotide excision repair
Fanconi anemia pathway
Ribosome
Coronavirus disease − COVID−19
TGF−beta signaling pathway
Oocyte meiosis
Progesterone−mediated oocyte maturation
Viral carcinogenesis
Human T−cell leukemia virus 1 infection
C1 (56)
C10 (93)
C11 (99)
C12 (137)
C13 (60)
C14 (49)
C17 (84)
C18 (46)
C2 (145)
C3 (114)
C4 (153)
C6 (57)
C8 (186)
C9 (124)
pvalue
2e−4
4e−4
6e−4
8e−4
GeneRatio
0.05
0.10
0.15
0.20
0.25

**Figure S5. Functional analysis of DEG clusters. (A-B)** The significant GO terms for biological processes **(A)** and KEGG pathways **(B)** for 20 DEG clusters.

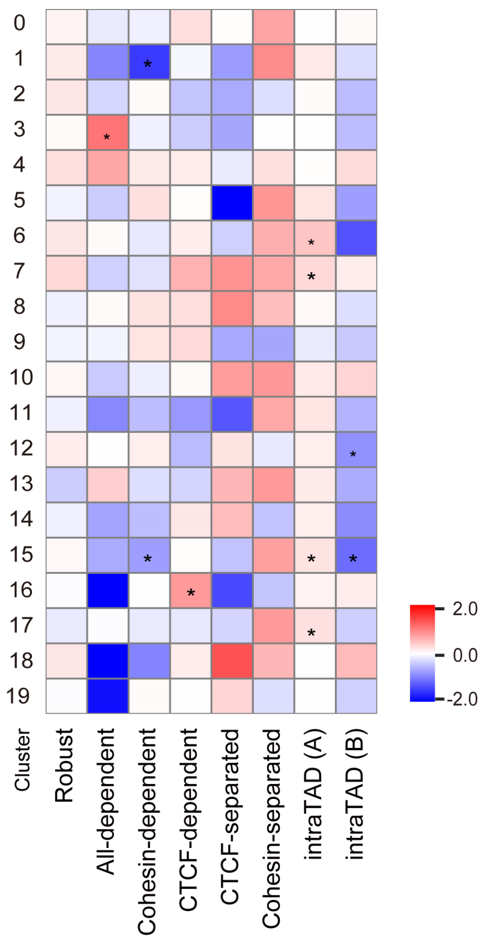


**Figure S6. siRNA effects on the transcriptome.** The relative enrichment of DEGs that overlap with the six types of boundaries and intraTAD regions for compartments A and B relative to all boundaries. Significance was determined by the permutation test (n = 1,000, *p < 0.01).

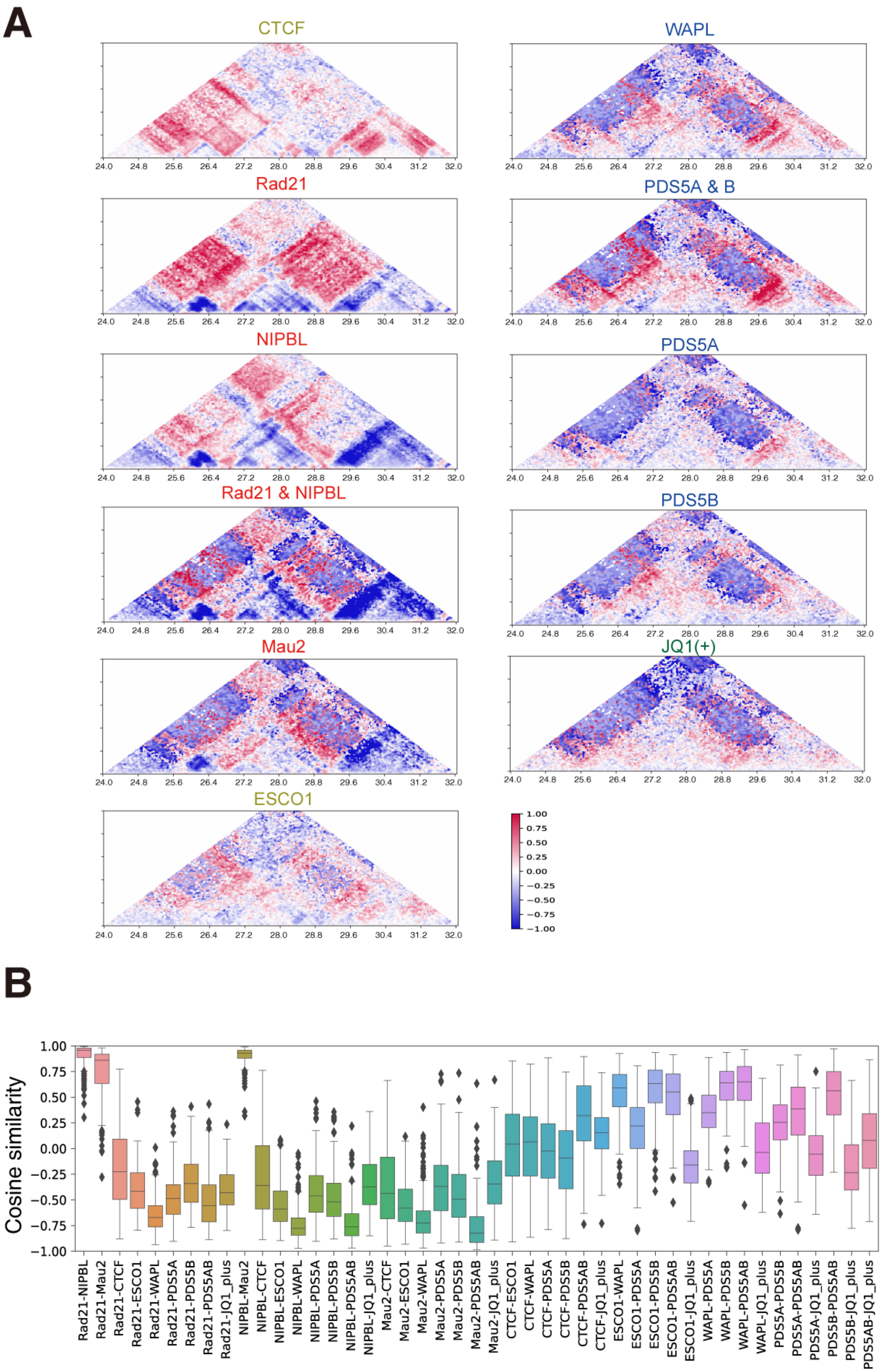

A
CTCF
WAPL
Rad21
PDS5A & B
NIPBL
PDS5A
Rad21 & NIPBL
PDS5B
Mau2
JQ1(+)
ESCO1
24.0 24.8 25.6 26.4 27.2 28.0 28.8 29.6 30.4 31.2 32.0
1.00 0.75 0.50 0.25 0.00 −0.25 −0.50 −0.75 −1.00
B
Cosine similarity
1.00 0.75 0.50 0.25 0.00 −0.25 −0.50 −0.75 −1.00
Rad21-NIPBL
Rad21-Mau2
Rad21-CTCF
Rad21-ESCO1
Rad21-WAPL
Rad21-PDS5A
Rad21-PDS5B
Rad21-PDS5AB
Rad21-JQ1_plus
NIPBL-Mau2
NIPBL-CTCF
NIPBL-ESCO1
NIPBL-WAPL
NIPBL-PDS5A
NIPBL-PDS5B
NIPBL-PDS5AB
NIPBL-JQ1_plus
Mau2-CTCF
Mau2-ESCO1
Mau2-WAPL
Mau2-PDS5A
Mau2-PDS5B
Mau2-PDS5AB
Mau2-JQ1_plus
CTCF-ESCO1
CTCF-WAPL
CTCF-PDS5A
CTCF-PDS5B
CTCF-PDS5AB
CTCF-JQ1_plus
ESCO1-WAPL
ESCO1-PDS5A
ESCO1-PDS5B
ESCO1-PDS5AB
ESCO1-JQ1_plus
WAPL-PDS5A
WAPL-PDS5B
WAPL-PDS5AB
WAPL-JQ1_plus
PDS5A-PDS5B
PDS5A-PDS5AB
PDS5A-JQ1_plus
PDS5B-PDS5AB
PDS5B-JQ1_plus
PDS5AB-JQ1_plus

**Figure S7. siRNA effects on long-range contact frequency. (A)** Relative enrichment of the interaction frequency (log scale) relative to control for all samples (same chromosomal region as shown in Figure 4A.) **(B)** The cosine similarity distribution of the relative frequency in all 241 differential DRF regions (~2 Mbp from the center of each region).

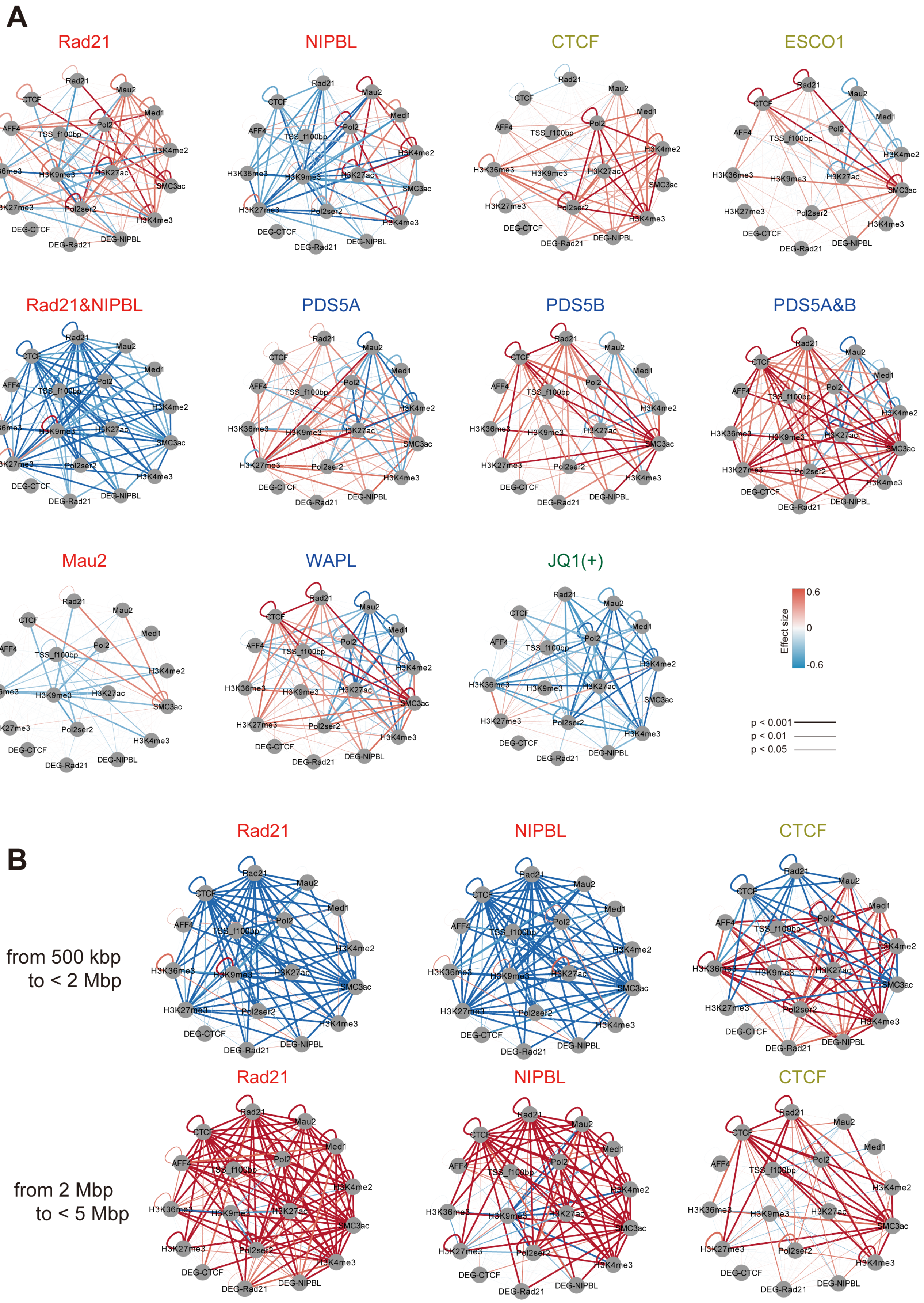

A
Rad21
NIPBL
CTCF
ESCO1
Rad21&NIPBL
PDS5A
PDS5B
PDS5A&B
Mau2
WAPL
JQ1(+)
Rad21
Mau2
CTCF
Med1
AFF4
Pol2
TSS_f100bp
H3K4me2
H3K36me3
H3K9me3
H3K27ac
SMC3ac
H3K27me3
Pol2ser2
H3K4me3
DEG-CTCF
DEG-Rad21
DEG-NIPBL
Effect size
0.6
0
-0.6
p < 0.001
p < 0.01
p < 0.05
B
from 500 kbp to < 2 Mbp
from 2 Mbp to < 5 Mbp
Rad21
NIPBL
CTCF

**Figure S8. SIMA analysis for all samples. (A)** SIMA analysis for all samples (distance, 500 kbp–5 Mbp). Color and width of edges correspond to the effect size and significance (Wilcoxon signed-rank test), respectively. **(B)** SIMA analysis for distances of 500 kbp–2 Mbp (top) and 2 Mbp–5 Mbp (bottom) for siRad21, siNIPBL and siCTCF.

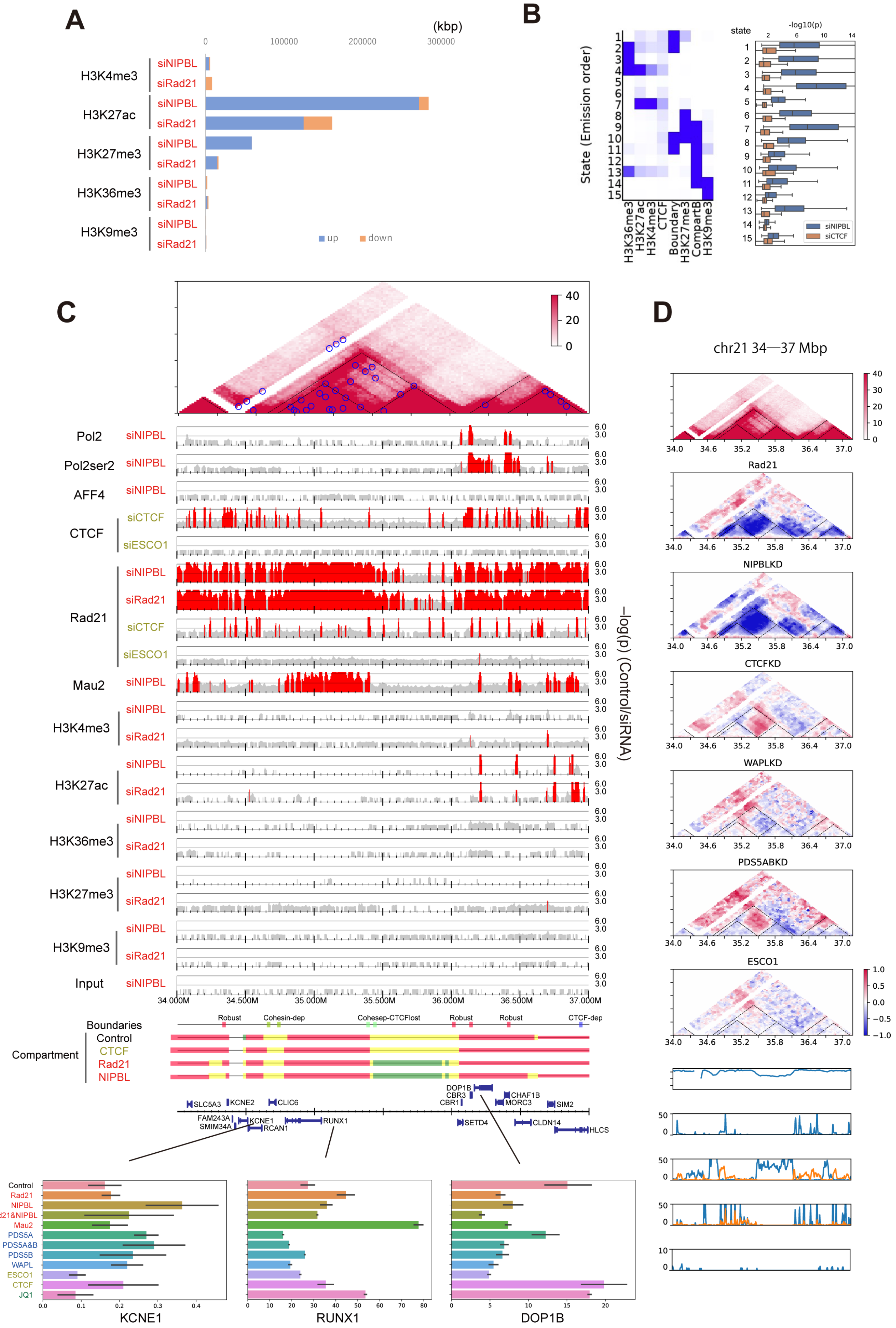

A
B
C
D
chr21 34—37 Mbp

**Figure S9. Correlation of depletion effects among epigenome, transcriptome and chromatin folding. (A)** The total width of genomic regions in which the histone marks were enriched or depleted for each siRNA target. **(B)** Left: Fifteen chromatin states using extended ChromHMM, which includes compartment PC1 and boundary features. Right: Distribution of $-\log_{10}(p)$ for the depletion effect of siNIPBL and siCTCF on Rad21 ChIP-seq in each state. **(C)** *RUNX1* locus (chromosome 21, 34–37 Mbp.) Top: Hi-C heatmap and the depletion effect distribution of ChIP-seq ($-\log_{10}(p)$, control/siRNA, 5-kbp bin.) Black dashed lines and blue circles indicate called TADs and loops, respectively. Middle: The four compartment types as indicated by the colored bars (red, StrongA; yellow, WeakA; green, WeakB; blue, StrongB). Bottom: Gene annotation and averaged gene expression level (TPM) for all siRNA targets. **(D)** Relative enrichment of interaction frequency and the $-\log_{10}(p)$ visualization of ChIP-seq data for the same region.

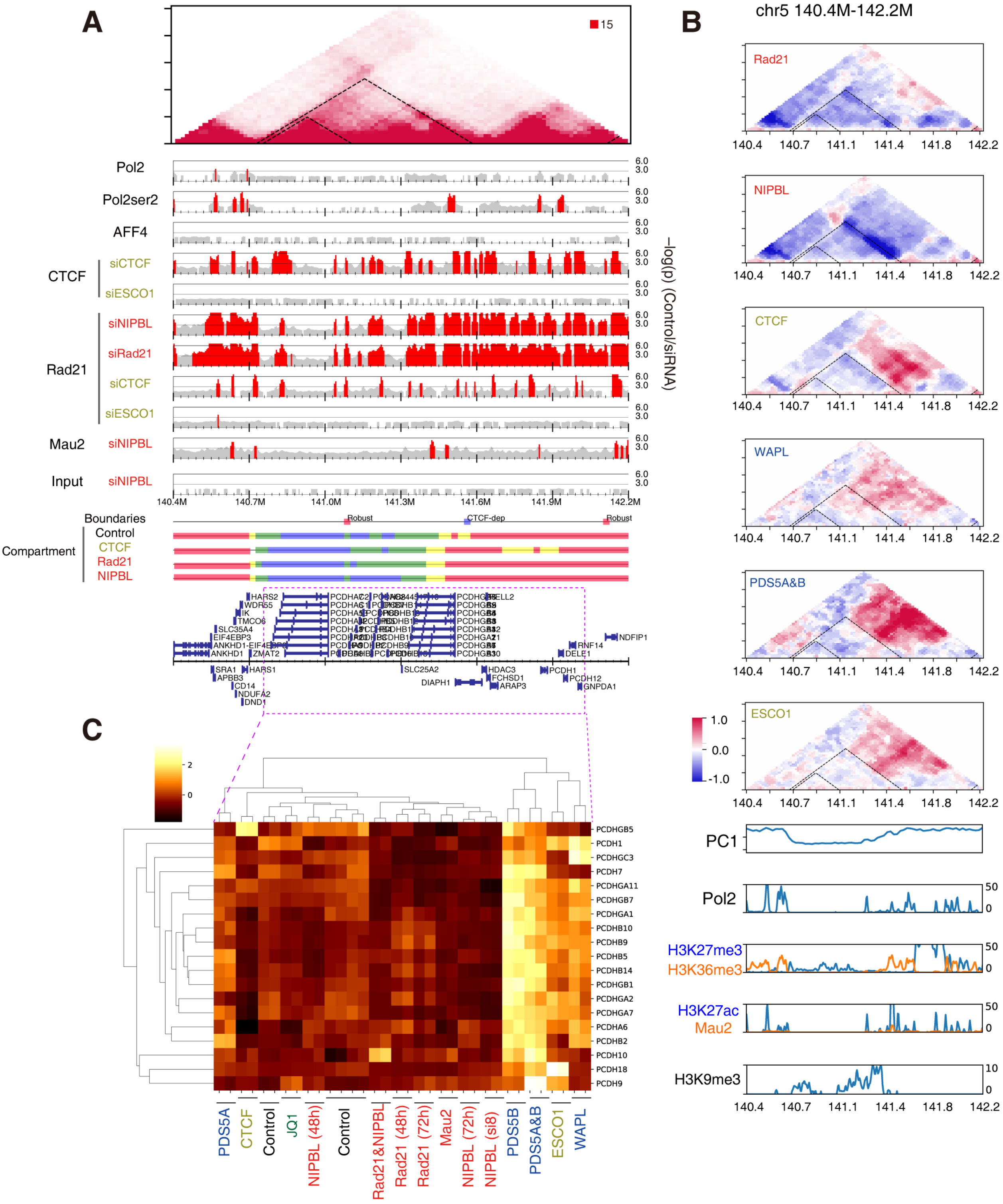


**Figure S10. InterTAD interactions and gene expression at *PCHD* genes (chromosome 5, 140.4–142.2 Mbp). (A)** The ChIP-seq distribution (5-kbp bin), analogous to Figure 7A. **(B)** Relative interaction frequency and ChIP read enrichment (–$\log_{10}$(p), ChIP/input, 5-kbp bin). **(C)** The expression level of *PCHD* genes that were identified as DEGs (TPM; Transcripts Per Kilobase Million, z-normalized).
